# The utility of whole-genome sequencing to identify likely transmission pairs for pathogens with slow and variable evolution

**DOI:** 10.1101/2024.05.06.592672

**Authors:** A. J. Wood, C. H. Benton, R. J. Delahay, G. Marion, E. Palkopoulou, C. M. Pooley, G. C. Smith, R. R. Kao

## Abstract

Pathogen whole-genome sequencing (WGS) has been used to track the transmission of infectious diseases in extraordinary detail, especially for pathogens that undergo fast and steady evolution, as is the case with many RNA viruses. However, for other pathogens evolution is less predictable, making interpretation of these data to inform our understanding of their epidemiology more challenging and the value of densely collected pathogen genome data uncertain. Here, we assess the utility of WGS for one such pathogen, in the “who-infected-whom” identification problem. We study samples from hosts (130 cattle, 111 badgers) with confirmed infection of *M. bovis* (causing bovine Tuberculosis), which has an estimated clock rate as slow as ∼0.1–1 variations per year. For each potential pathway between hosts, we calculate the relative likelihood that such a transmission event occurred. This is informed by an epidemiological model of transmission, and host life history data. By including WGS data, we shrink the number of plausible pathways significantly, relative to those deemed likely on the basis of life history data alone. Despite our uncertainty relating to the evolution of *M. bovis*, the WGS data are therefore a valuable adjunct to epidemiological investigations, especially for wildlife species whose life history data are sparse.

## 1 Introduction

Pathogen whole-genome sequencing (WGS) is an invaluable tool for understanding transmission patterns in infectious diseases [1–4]. Underpinning such use of pathogen WGS data is the relationship between the rates at which evolutionary processes occur (the “molecular clock” [5]), and the timescale of epidemiological processes of interest (e.g. the generation time between infection events). If many events are observed and occur at a consistent rate, this allows for precise data to inform our understanding of the epidemiology. However, for many bacterial pathogens this evolution can be highly variable, and slow enough that the timescale for a single variation is comparable to or longer than the generation time of the disease itself [6, 7]. Here, it is less clear how useful WGS data may be for epidemiological studies [8].

An important example of a slowly evolving pathogen for which considerable WGS data has been gathered is *Mycobacterium bovis* (*M. bovis*), a member of the *Mycobacterium tuberculosis complex* (MTBC) and the causative agent of bovine Tuberculosis (bTB) in many mammal species [9]. The *M. bovis* genome comprises ∼4.3 ×10^7^ base pairs [10], and has an estimated evolutionary rate of between 0.1 − 1 variations per year [11–15]. However many estimates from the literature are non-overlapping, consistent with a highly variable evolutionary rate.

Bovine Tuberculosis is a chronic zoonotic disease with considerable animal health and economic impacts in many countries including in the United Kingdom and Ireland [16–18]. Control of bTB is a complex challenge in part due to a low sensitivity in the standard skin test for disease in cattle [19], and in several countries the presence of additional susceptible wildlife species capable of cross-species transmission (e.g. European badgers (*Meles meles*) in the UK and Ireland, wild boars (*Sus scrofa*) in continental Europe, white-tailed deer (*Odocoileus virginianus*) and elk (*Cervus canadensis*) in the USA and Australian brushtail possums (*Trichosurus vulpecula*) in New Zealand).

Studies using *M. bovis* WGS data so far mainly focus on inference of *phylogenetic* trees and the evolutionary history of the pathogen. This has been used to characterise clusters of transmission [20–24], and infer rates of between-species transmission from when the sample species changes between nodes in the phylogenetic tree [12,15,20,25–30]. This is a first step towards building more complete *transmission trees*, and inference of *who-infected-whom*. However, for slower-evolving pathogens different host samples may have very similar or even identical sequences entirely, making it more diffcult to resolve specific infector-infectee pairs [31]. There are a family of approaches for transmission tree reconstruction that combine evolutionary and epidemiological processes [32], including Bayesian inference methods which have been applied to MTBC pathogens [33–36]. Reconstructions on simulated data (where the transmission tree is known) indicate that tree accuracy is sensitive to sampling density [37], and for *M. bovis* in particular, suffers when wildlife species are undersampled [38]. These are key challenges especially for multi-species disease systems such as bTB, where understanding the role of different species is central to effective disease control [39].

Towards this, then, here we assess the utility of *M. bovis* WGS data in informing *who-infected-whom*. We study WGS samples taken from cows and badgers with confirmed *M. bovis* infection in the vicinity of a long-term study of infection in badgers (Woodchester Park) [40]. These data are detailed but low-density, and have significant sample biases which make a full transmission tree inference unsuitable. Instead, then, starting with *all* possible pathways, we assess how the sample data can be used to directly *rule out* pathways, leaving only those that are supported by the sample data. Our interest is in how WGS data can be used to further discriminate between different pathways, by either corroborating existing data, or contradicting them (ruling out pathways that otherwise appear plausible, or boosting pathways where the host data do not support a transmission, but their WGS data do).

For each potential transmission pathway between sampled hosts, using an underlying model for *M. bovis* evolution and transmission we calculate the *relative likelihood* that such a transmission occurred (defined here as the size of space of model outcomes that are consistent with all sample data on those hosts). This is informed by the host sampling times, life history data (time of birth and death, location and movement history), and rates associated with the epidemiology of bTB. In a second set of calculations we include the number of single nucleotide polymorphisms (SNPs) identified in the pathogen sequences relative to a reference sequence. Any differences found between the two approaches over different pathways indicate the additional discriminatory power of the WGS data. We hypothesise that, despite the slow and variable evolution that *M. bovis* exhibits, WGS will be a powerful complementary source of information for inferring transmission pathways, when used alongside existing life history data.

## 2. Results

We used cattle WGS samples where hosts had a known birth and death date, and movement history between holdings, and badger WGS samples where hosts had a known sample location (their lifespan is estimated from their capture history, see Section 4). We also removed samples from before the year 2000 due to the unreliability of cattle tracing data before this period, as well as repeat samples from the same host (taking only the first sample). This left *M. bovis* samples from 111 badgers and 130 cattle, spanning between 2000 and 2020 (spatial and temporal distribution shown in Supplementary Material A, Fig S1). Most of the cattle samples were from 2013–2020. We note this represents a low density of sampling relative to all known cases in badgers and cattle in the vicinity of the study area and to all likely infections; recorded and not recorded (see Supplementary Material A, Fig S2 for comparative plots including all cases including those not sequenced).

The 241 samples gives a maximum of 241 × (241 − 1) = 57840 potential transmission pathways to assess (each host pair, each direction). We removed those deemed impossible due to the hosts not being alive at the same time, and cattle pairs that according to the cattle movement data had not been in the same holding at the same time. This left 12844 possible direct pathways. For pairs of cows that did not necessarily interact directly, but where each shared a holding with one or more cows that may have acted as an intermediary host, we calculate the relative likelihood of transmission via those intermediaries. Of the 130 cattle (16670 potential cattle-cattle pathways), we found 56 pathways with direct interaction, and 247 pathways via potential intermediate hosts. Longer transmission chains with additional intermediaries were not considered but would be considered as having diminishing added value with increased chain length (see Supplementary Material B, Table S1 for a summary of transmission pathways considered and not considered in this work).

For all transmission pathways remaining, we then calculated the relative likelihood of such a transmission having occurred, via a numerical integration. This calculation is outlined in Section 4.3, with full integral expressions (which are not analytically solvable) presented in Supplementary Material C. Briefly here, we model a 2-body susceptible-exposed-infectious process, where substitutions occur on the *M. bovis* genome via a Poisson process. We define here the *relative likelihood* of transmission as the size of the state space of model trajectories under our model and a given choice of model parameters, that would be consistent with the observed data and genetic distance between samples.

### 2.1 Relative likelihood over all transmission pathways

Fig 1 illustrates relative transmission likelihoods across all hosts in the data, with hosts ordered by time of sampling. For transmission between cattle pairs via an intermediary host, these pathways are illustrated in Supplementary Material, Fig S6. Line thickness indicates the relative transmission likelihood. For each host pair H_1_, H_2_, the line is for the pathway between those hosts with the highest relative likelihood (H_1_ →H_2_/H_2_ →H_1_). For the variability in relative likelihood in these datasets, many pathways result in line thickness that is too fine to be resolvable.

**Figure 1.**
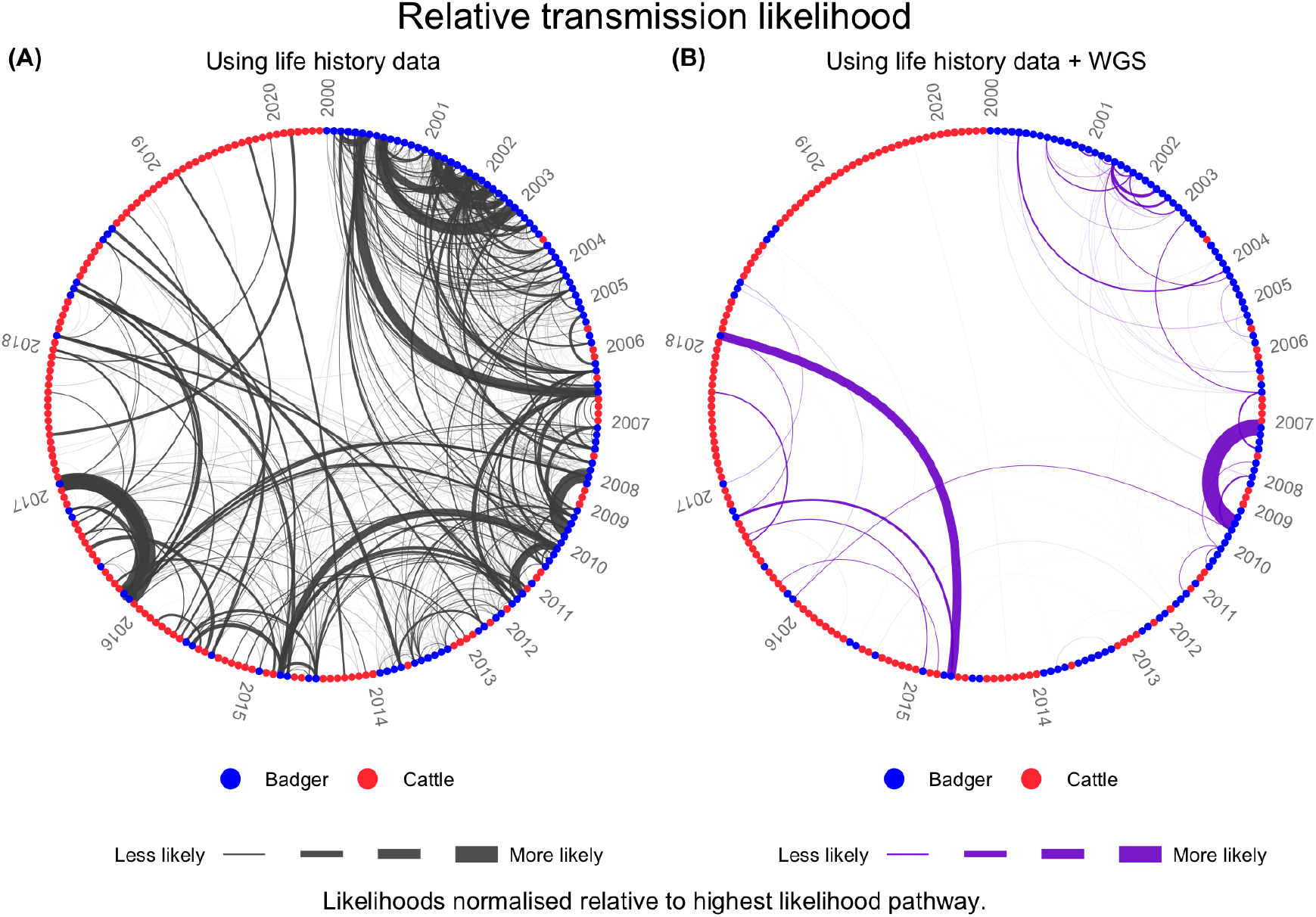
Relative direct transmission likelihoods between all hosts, evaluated on the basis of (**A**) life history data, (**B**) life history data and WGS data. The width of the connecting line is the relative likelihood. For each pair of hosts H_1_, H_2_ (each host being a point, the colour denoting the species), the relative likelihood is evaluated in both directions 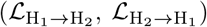, and we show here the pathway with the higher relative likelihood. Samples are time-ordered in a clockwise arrangement, with the earliest samples at “twelve o’clock”. The relative likelihoods in each set are scaled relative to the pathway with the highest relative likelihood; on this scale many pathways result in line thickness too fine to be resolvable.

Over both sets of relative likelihoods there is a clear temporal pattern, with higher relative likelihoods for host pairs sampled closer together in time (see Supplementary Material A, Fig S1 for the temporal distribution of samples). Higher relative likelihood pathways are mainly same-species (see Supplementary Material D, Figs S4, S5 for a breakdown by species pair).

The lines between hosts representing the relative likelihoods are characteristically sparser in Fig 1B than in Fig 1A, indicating a larger spread. A broader distribution of relative likelihoods (on a log scale) indicates stronger discrimination between different pathways. The standard deviation of the relative likelihoods over all pathways quantifies the degree to which the inclusion of WGS data has further discriminated between different pathways. Excluding those with likelihood zero, the standard deviation of relative likelihoods evaluated on the basis of life history data is 26.4 (powers of 10), and with life history and WGS data combined is 46.7 (see Supplementary Material D, Fig S7 for full distributions).

### 2.2 Comparing the discriminatory power of WGS over different species

Fig 2A shows the distribution of relative likelihoods over all transmission pathways, on the inclusion of WGS data, broken down by pathway type. The specific value of the relative likelihood for all transmission pathways decreases with the inclusion of WGS data. The degree of *y*-axis dispersion then illustrates that the WGS data further discriminates between transmission pathways with similar relative likelihoods, when evaluated on the basis of life history data alone.

**Figure 2.**
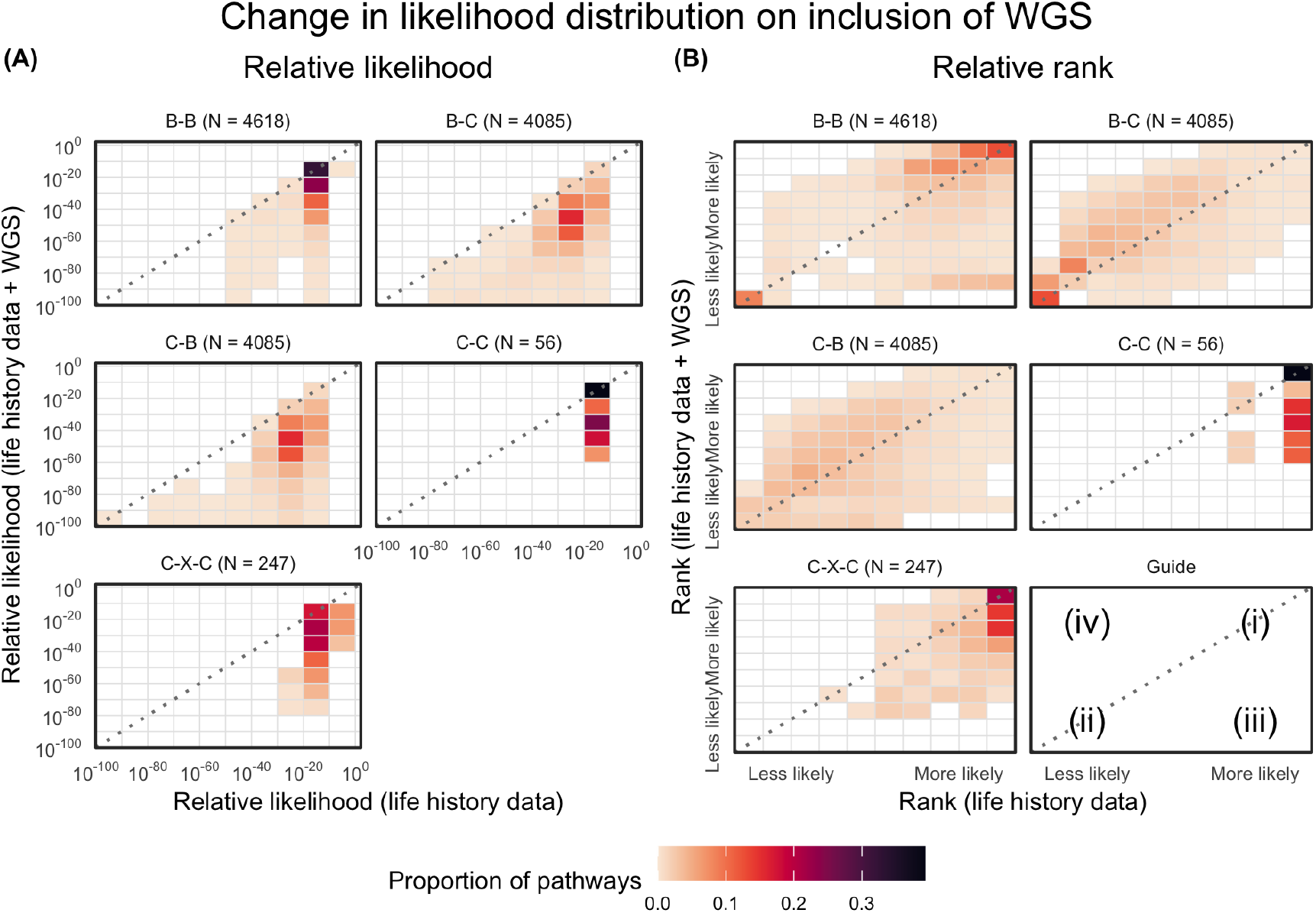
(**A**) Density plots showing the distribution of relative likelihoods for individual transmission pathways, on the basis of life history data (*x*-axis), and life history data and WGS data (*y*-axis). Darker squares indicate a higher proportion of pathways. Each subplot covers a different species pair, noting significantly different total numbers in each. We exclude pathways deemed impossible from life history data alone. (**B**) The *rank* order of all pathways from highest to lowest relative likelihood, again on the basis of life history data (*x*-axis), and the inclusion of WGS data (*y*-axis). Points on the diagonal indicate pathways where the additional WGS data broadly corroborates the relative likelihood evaluated on the basis of host life history data alone. Regions away from the diagonal are where the WGS data contradict the evidence evaluated on the basis of life history data alone; these are either deemed to now be relatively less likely (bottom right), or more likely (top left). The position of each of these pathways places them into broad categories *(i)*–*(iv)* in Table 1.

To quantify this discriminatory power, in Fig 2B we have compared the *ranks* of transmission pathways in order of relative likelihood. This allows us to classify different transmission pathways in to four broad categories (see Table 1), to indicate whether the transmission pathway is supported, and also whether the WGS and life history data corroborate. Pathways that deviate from the diagonal in Fig 2B fall in to categories *(iii)* and *(iv)*, where WGS contradicts the life history data (*(iii)*: deviating bottom-right, *(iv)* deviating top-left).

**Table 1:** A broad categorisation of transmission pathways, considering the relative likelihood evaluated on the basis of life history data alone, and that evaluated on the inclusion of WGS data.

| Label | Description | Examples |
| --- | --- | --- |
| (i) | The pathway is relatively *likely* on the basis of both life history data and WGS data. The transmission is supported by both data sources. | Supp. Mat., S3 |
| (ii) | The pathway is relatively *unlikely* on the basis of life history data and WGS data. The transmission pathway can be reasonably ruled out. | Supp. Mat., Table S4 |
| (iii) | The pathway is relatively *likely* on the basis of life history data but WGS is *contradictory* to this. The transmission can be reasonably ruled out on the basis of WGS data. | Supp. Mat., Table S5 |
| (iv) | The pathway is relatively *unlikely* on the basis of life history data but WGS is *contradictory* to this. Despite the life history data being unsupportive, the WGS signature merits further investigation (e.g. seeking a missing intermediary, or considering a transmission scenario outwith prior assumptions of the epidemiology of bTB). | Supp. Mat., Table S6 |

The two cattle-cattle pathways with the highest relative likelihood from life history data are between hosts labelled C.054 → C.034 (rank 65 out of 13091 pathways, including pathways via an intermediary) and C.104 → C.080 (rank 66). These pairs each spent extended time together in the same holding. Taking the WGS data, the genetic distance between the samples for C.054 and C.034 was 4 (C.054 had 1 unique SNP relative to a reference sample not seen in C.034, which in turn had 3 unique SNPs not seen in C.054), and the relative likelihood with these data remains high compared to other pathways (rank 22, category *(i)* in Table 1). Conversely, the genetic distance between the samples for C.104 and C.080 was 33 (with 18 and 15 unique SNPs respectively). Such a divergence between infector and infectee sequences is extremely improbable given the time between observation and possible infection (see Eq (10)) ruling it out as a potential transmission pathway (rank 4575, category *(iii)* in Table 1). The relative likelihoods of these pathways are illustrated in Supplementary Information E, Fig S8A, amongst similar pathways. We also include in Supplementary Material E, Fig S8B, C, a set of pathways infecting two specific hosts: a badger labelled B.104 (the infectee in the transmission pathway with highest relative likelihood on the basis of life history data), and cow C.085 (the cow with the most interactions with other sampled cattle, direct and via-intermediary). This mimics an epidemiological investigation (asking who infected a particular host). For both, the inclusion of WGS provides a significant degree of discrimination, with the majority of pathways fitting category *(iii)* in Table 1. For C.085 the WGS data identify the most likely source of infection amongst sampled hosts to be a badger, ∼B.088. This badger was sampled ~1 km from a holding that the cow had spent time in, and the WGS samples had a genetic distance of 2 SNPs.

Further examples of specific transmission pathways and relative likelihoods are given in Supplementary Material, Tables S3–S6, with details of the host life histories, and genetic distances amongst *M. bovis* sequences.

#### Sensitivity to parameter choices

The rank ordering of transmission pathways here are from median relative likelihoods over many parameter draws. The broad rank ordering of pathways remains consistent when looking at individual parameter draws (distributions given in Supplementary Material, Table S2). The median “range of ranks” for pathways under different draws was 362 (with 11077 total having likelihood greater than zero, see Supplementary Material, Fig S10A). In other words, more likely pathways are consistently more likely, and less likely pathways consistently less likely. However, this variability emphasises the need to consider these relative likelihoods with their broad confidence intervals in mind (as illustrated in e.g. Supplementary Material, Fig S8).

We also assess the sensitivity of the relative (log) likelihood to changing each of the variables in the model via a *Boruta* analysis. This is a standard machine learning method applied to multivariable data that assesses how “important” different variables are in explaining an outcome (the relative likelihood). This is based on how much accuracy a statistical model *loses* if a variable was effectively removed; more important variables see greater losses. This showed (Supplementary Material, Fig S10B) that the relative likelihood is initially most sensitive to changes in the infection rate *β*_0_, and to the rate of transition to infectiousness *σ* and time to test-sensitivity Ω to a lesser degree. However on the inclusion of WGS data, the substitution rate *μ* is overwhelmingly the most important variable; the relative likelihood is most sensitive to changes in *μ* as opposed to other variables.

## 3 Discussion

Pathogen whole genome sequencing data, coupled with an understanding of that pathogen’s evolution, can be used to estimate a time to a most recent common ancestor. This in turn may inform the relative likelihood that one host infected another. Used in conjunction with other host data, WGS provides clear utility for inferring transmission pathways when pathogen evolution is steady and predictable, and a reasonable number of substitutions are expected to occur between infection and onward transmission. The aim of this work was to assess the utility of WGS in a far more challenging context applicable to many important pathogens: when evolution is sufficiently slow that the substitution rate is itself difficult to estimate. This applies to wide range of bacterial pathogens, including mycobacteria such as *M. bovis*, which evolves at an estimated rate as low as one substitution on the genome every 10 years. In this instance, the number of variations between isolates from different hosts can be zero. The “who-infected-whom” identification problem is especially pressing for *M. bovis* control in multi-host systems where infection may be transmitted amongst susceptible domestic and wild species.

We described a method for calculating the *relative likelihood* of transmission between specific hosts in a sample of cattle and badgers with confirmed *M. bovis* infection. We find that a relatively large number of plausible transmission pathways can be identified on the basis of life history data alone (when the hosts were alive, and where). However, the inclusion of data on genetic distances between *M. bovis* isolates permitted the number of plausible pathways to be reduced considerably, ruling out those where the genetic distance between samples would be highly improbable based on the evolutionary characteristics of the pathogen. This illustrates the utility of WGS in identifying a subset of more likely transmission pathways, despite uncertainty over the evolutionary rate of the pathogen (using a wide range of *μ*∈ U(0.1, 0.9) substitutions/genome/year).

The manner in which WGS was “useful” in our analyses depended on the specific transmission pathway, as categorised in Table 1. First, WGS may *corroborate* the relative likelihood evaluated on the basis of life histories, either as *(i)* more likely or *(ii)* less likely. Second, in some instances *(iii)* the pathway is deemed likely given life history data, but the genetic distance between pathogen samples is highly improbable, ruling out that pathway. These are important for “who-infected-whom” investigations in further pruning pathways that appear plausible based on all other data. The final category *(iv)* is where an unlikely pathway emerges as comparatively *more* likely on being informed by WGS. An example of this may be hosts that were never in close proximity to one another, but the genetic distance is consistent with a direct transmission having occurred. These pathways merit further investigation. For example, there may have been an unobserved intermediary(s) in a longer transmission chain connecting the two hosts, in which case the life history data can be used to identify where intermediaries may have been. Examples include additional cattle in the movement network, or wild animals at intermediate physical locations. We could also interpret these results as a challenge to the assumptions we have made about the epidemiology of bTB; that is, a mechanism of transmission not captured by our model assumptions (e.g. fomites, rare long-distance badger movements [41]).

The transmission relative likelihoods are evaluated directly by integrating over the different epidemiological stages (exposure, infectiousness, onward infection), bounded by known delays in test sensitivity and life history data of the hosts. The epidemiology of bTB and evolution of *M. bovis* are embedded within these formulae as functional expressions, which may be generalised to more sophisticated epidemiology (e.g. a more elaborate compartmental model) and models of pathogen evolution. The method presented here accounts for transmission between cattle via a possible single intermediary that has not been sampled, and this could be generalised to longer chains of transmission. An estimate of the potential number of intermediary hosts could also be important in defining the probability of a wildlife host (in areas where wildlife are not sampled) being involved, given information on the testing frequency in cattle.

### Method limitations and assumptions

This approach is by construction highly sensitive to the underlying data and model assumptions. The main limitation is how accurately the model and data capture the true movements and interactions of the hosts, and the epidemiology of bTB. Owing to the slow progression of disease and variability in disease severity amongst hosts, many of the parameters used in this work are highly uncertain.

On the evolution of the *M. bovis* bacterium, we model substitutions as occurring randomly and independently at some prescribed rate (Eq (9)). While the range of substitution rates reflects that estimated in the literature [11–15], these estimates are often non-overlapping and suggest that the “true” substitution rate may be non-regular in time, and also vary depend on the pathogenesis of *M. bovis* and the immunological response to infection in specific hosts, as well as the specific lineage of *M. bovis*. Inevitably, the collection of more WGS data will allow us to build a more precise understanding of the evolutionary process and these systematic variations, and in turn improve the value of those data in such epidemiological investigations.

A limitation specific to the cattle data is that while we identify “same holding, same time” instances for cattle pairs, holdings themselves may be geographically fragmented meaning some cattle on the same holding may not have had any opportunity to interact. In addition we may miss important interactions such as those “over-the-fence” between adjacent holdings, or transmission “between” two hosts that had never interacted directly, via fomites (such as a common piece of equipment, or contamination of shared grazing land).

Our model choice (a two-body susceptible-exposed-infected model) is relatively simple and, before considering lifespan and location, places few constraints as to the types of transmission permitted. The same approach used here could be applied with a more complex disease model that considered e.g. details of the environment and natural boundaries, social structures and roaming behaviour. Such an approach would mainly rule out more pathways on top of those done so here, rather than returning an entirely different set of plausible/implausible pathways.

Our model uncertainty is greatest in the badgers, as these wild animals do not have precisely tracked locations. The badger data we use here are exceptional, as the social structure and hence the resident territories of badgers in Woodchester Park are well documented. We estimated a distance-dependency on the infection rate from the difference between within and between-social group infection rates (Eq (7)). The main effect of changing this distance dependency is to systematically suppress/amplify pathways at higher distances (see Supplementary Material F, Fig S9), but any distance-dependency on infection probability between two hosts will vary based on factors including but not limited to hosts’ tendency to move and the influence of the distribution of suitable habitat and physical boundaries to movement, and the nature of their interactions with others (e.g. badgers biting one another in territorial disputes).

The infection rates used in this work can be refined further, however sensitivity analysis from Section 2 does indicate that the relative likelihood is overwhelmingly more sensitive to a tuning of the substitution rate. Thus though we would expect some changes to individual rankings, the overall conclusions as to the discriminatory value of WGS even with such low substitution rates, are unlikely to change.

Finally, we do not consider historical negative tests from either species, which may inform lower bounds on infection time. We chose not to include this due to the poor sensitivity of the diagnostic tests for bTB [19,42]; as all sampled hosts in these data recorded a positive test eventually, previous tests may in fact have been false negatives. We were then not confident in including a hard cutoff at, say, the most recent negative test, however these data could be used in a more careful manner in the future, if considering test sensitivity and overall disease prevalence.

### Sample coverage

For the data used here, the sample coverage relative to all likely infections within the study area in the period studied is very low. We analysed 130 cattle between 2000–2020, which is only a small subset of the 5852 cattle in total found with *M. bovis* infection (either a skin test reactor, or visible lesions found at slaughter) within a 10 km radius of the Woodchester Park study area (per the SAM test database). Similarly, over 715 badgers were found with confirmed infection (either by culture or serological testing) during this period in Woodchester Park alone, inclusive of the 111 sequences studied (Supplementary Material, Fig S2). With the compounding effect of not all infections being detected by the tests available, this makes it unlikely that many “true” transmission pairs are fully characterised by these data, particularly between-species due to strong spatial and temporal biases in host sampling (see Supplementary Material, Fig S1). For these reasons we hesitate to make strong claims about “direction of transmission” from this analysis (as compared with analyses that build the broader phylogenetic tree [12, 15, 20]). Rather, this work indicates that WGS is highly effective for *ruling out* transmission pathways across sampled hosts, including instances when transmission appeared likely on the basis of other information (Supplementary Material, Fig S8B). This approach provides a simple method for assessing when an infection source has been missed (i.e. when for a specific host, *no* plausible infection source is found). A simple application would be to infer a wildlife source for a herd breakdown, where no suitable cattle source was found after dense cattle sampling.

### Specific utility of WGS for disease transmission in wildlife

Knowledge of cattle movements alone allowed us to eliminate the vast majority of potential transmission pathways amongst cattle (notwithstanding limitations of these data as described above), though WGS was effective at discriminating between remaining pathways (e.g., Supplementary Material, Figs S5, S6, S8C). Conversely, without knowing such detailed movement data for individual wild badgers (even though the data used were exceptional for a wildlife species), we had to consider all pathways involving hosts with overlapping lifespans. For these pathways WGS was especially effective and often contradicted the relative likelihood evaluated on the basis of only life history data (e.g. Fig 2B). This indicates that WGS has particular value for interactions involving wildlife species despite the variability in pathogen evolution, as other data on movements and propensity to transmit disease are likely to be even more limited. This is especially relevant to *M. bovis*, where a principle challenge in disease control in several countries is understanding the role of the wildlife reservoir in sustaining infection in domesticated species. In a separate case of cattle infection in the Low Risk Area of England where badger culling had taken place, there was sufficient WGS data on wildlife to potentially determine the field in which cattle grazed and were infected, as we had more detailed wildlife data and could potentially exclude one of the two areas where they had grazed. Such detailed analysis could only be determined from robust WGS data in both species.

## 4 Data and methods

### 4.1 Data

This study uses life history data on badgers and cattle located in and around Woodchester Park, a Site of Special Scientific Interest in south-west England. This lies within the bTB “high–risk” zone, which covers all of south-west England and some counties beyond. In this zone cattle testing is generally mandated every 6 months [43].

#### WGS samples

The WGS dataset contains *M. bovis* sequences with metadata on host species, sample date, and for a subset of the hosts a grid-reference location. WGS samples here are from between 2000 and 2020. In this period the primary method of bacterial characterisation in GB cattle was not WGS, but rather spoligotyping and variable number tandem repeat typing (both of which sample only short regions of the genome) [44, 45], with surveillance in wildlife beyond Woodchester Park extremely limited. The sequences dataset used here had been aligned to the reference genome AF2122/97 [46].

#### Woodchester Park dataset

The badger population in Woodchester Park has been intensively studied through a regular programme of capture-test-release since the 1970’s [40, 47, 48]. The study archives the life histories of captured badgers, including bTB test results, social group affiliation and capture locations. Clinical samples are collected from each captured badger for the purposes of serological testing and microbiological culture to determine their infection status. All captured animals are re-released unharmed following examination and sampling, regardless of their bTB status. Tests are conducted via culture or serological testing.

Exact times of birth and death of the badgers are not known. Most badgers are captured more than once, implying that almost all badgers are represented in the study data. Some badgers are first captured as cubs, whereas others may be found dead (often after a road traffic collision).

#### Cattle movements

The Cattle Tracing System (CTS, version dated 1 November 2020) archives the movements of all individual cattle in Great Britain, and the grid-reference locations of holdings [49]. From these data we associate with each WGS sample a time of birth and death, and movement history of the host cow, and the movement history of all cattle that had shared a holding with that cow at some point in its lifetime.

#### Cattle bTB test results

The SAM database (version containing entries up to 2021) contains data on cattle bTB test results, and herd breakdowns [50]. We extract from SAM the eartags of all cattle that had registered a positive bTB test, regardless of whether they were sequenced.

### 4.2 Data preparation

For each host, we require in addition to their WGS sample:

- time of sampling;
- time of birth and death (estimated for badgers based on observation times);
- location (observation location for badgers, full movement history for cattle).

#### Badgers

For each badger isolate, the corresponding isolate ID is linked to a badger in the Woodchester Park dataset. Captured badgers are released after examination and sampling regardless of infection status, so it is possible that they remain alive for several years after the sampling event. For badgers first captured as a cub, and later recorded as dead, we fix these dates as the times of birth and death respectively. For badgers first observed as an adult, or never observed dead, we use an estimated lifespan. For all badgers the birth date can be estimated as being between January-February, as per the modal birth time reported by Neal and Cheeseman [51]. For badgers first observed as a cub, we take the birth date to be February 1st of the year of first capture. For badgers first observed as an adult, we instead take the birth date as February 1st of the previous. For the death date, we either take the date of observation if the badger was found dead, or six months after the date of last capture if never found dead. The badger’s location is taken as the location of the badger at the time the sample was taken.

#### Cattle

For each cattle isolate, the corresponding isolate ID is linked to a cow in the CTS dataset (table `tblAnimal`), usually via an eartag. For hosts without eartags, we instead use the geographic location and time of the sample to identify from SAM potential cattle reactors. If a unique match is found, that eartag is used.

Cattle reactors are generally slaughtered very shortly after detection, thus the sample time is usually very shortly before the time of death.

### 4.3 Evaluation of relative transmission likelihoods

The relative likelihood that a transmission occurred from one given host to another is expressed as an integral over the ensemble of scenarios where one host infects another, constrained by:

- the *epidemiology* of bTB and the timescales in the epidemiological processes;
- the known *lifespan* of the badgers and cattle observed;
- the known *locations* of the badgers and cattle relative to one another;
- the *observation times* when samples were taken;
- the *evolutionary* process of *M. bovis* in the generation of substitutions on the genome.

Here we describe how each of these model assumptions and data inform the relative likelihood. The exact integral expressions for the relative likelihood are presented in Supplementary Material, Section C. Parameter ranges are summarised in Supplementary Material, Table S2.

#### 4.3.1 Epidemiology of *M. bovis*

We model bTB as a three-stage SEI (Susceptible, Exposed, Infectious) process (Fig 3). (I)nfectious hosts expose (S)usceptible hosts at rate *β*, denoting the time of exposure of host X as λ_X_. (E)xposed individuals transition to the infectious stage at rate *σ*, with the time of transition for host X denoted *ξ*_X_. The times λ and *ξ* are not generally observable.

**Figure 3.**
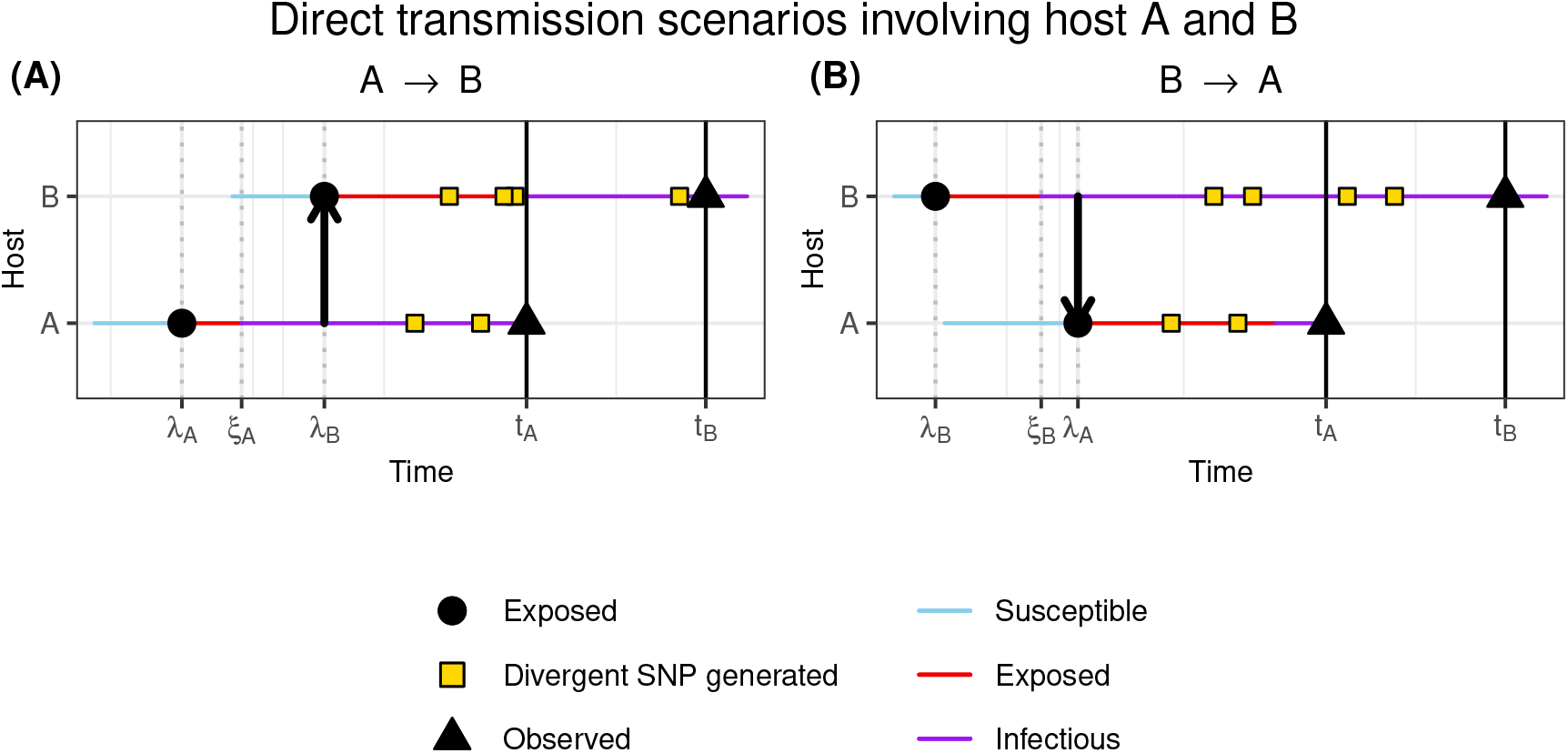
For two hosts A and B that are sampled at times *t*_A_ and *t*_B_ respectively, two transmission pathways that would be consistent with an observed genetic distance of 6 SNPs, with A having 2 SNPs not seen in B, and B with 4 SNPs not seen in A, both with respect to a reference genome. Hosts are born susceptible (blue), exposed to infection at times denoted λ (red), and become infectious at *ξ* (purple). The generation of SNPs on the pathogen genome inside each host are denoted with yellow squares. (**A**): A exposes B to infection. (**B**): B exposes A to infection.

Consider a system with two hosts A and B where at time λ_A_ A is exposed, with B susceptible. In an isolated system, A first transitions to infectiousness at some later time *ξ*_A_ ≥ λ_A_, after which A exposes B at some later time λ_B_ ≥ *ξ*_A_ (Fig 3A).

The time *ξ*_A_ depends on λ_A_; if A is exposed at λ_A_, the probability that A is not yet infectious by time *ξ* is

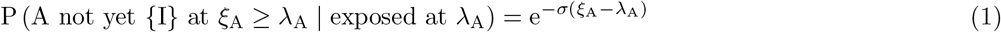

thus the distribution of transition times *ξ*_A_ is

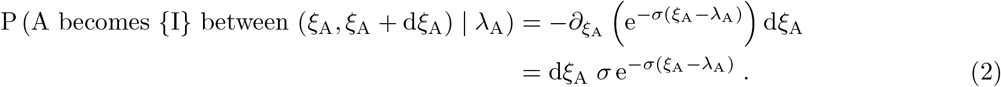

In a similar way, the time λ_B_ is conditioned on *ξ*_A_, and is distributed as:

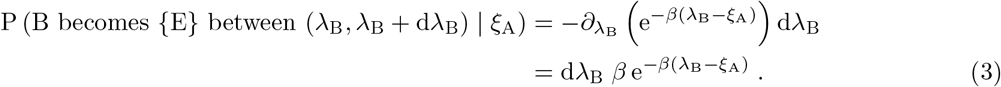

To evaluate the relative likelihood of transmission from one host to another, we integrate over all possible times of exposure *ξ* and transition λ within the system, constrained by the lifespans of the animals and times of observation.

#### 4.3.2 Test sensitivity

We assume that hosts are not immediately test-sensitive after exposure; that is, if tested, they would report a false negative. Rather, they only become test-sensitive after a period Ω. In our model, we take this time to be independent of the time of transition to *infectiousness*. We incorporate this with a term *W*_X_

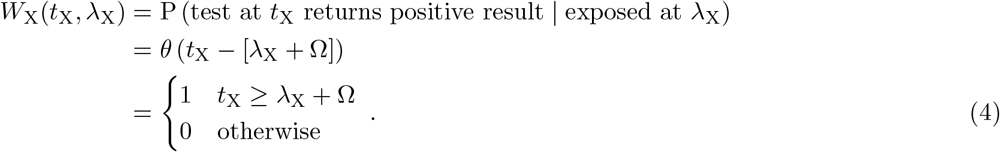

*θ* is a Heaviside step function; *θ*(*x*) = 1 if *x* ≥ 0, and 0 otherwise. This excludes scenarios where a host tests positive very shortly after being exposed to infection.

The time to test-sensitivity is another highly uncertain quantity (see e.g. [52], where the period was estimated as either within (0, 7.7) or (24, 517) days, dependent on model choice). We use the univariate range Ω ∈ U(0, 84) d based on a challenge study taking a skin test 12 weeks postexposure [53].

#### 4.3.3 Lifespan of hosts

The relative likelihood of transmission between a pair of hosts also depends on the overlap of their lifespans. If a host X was born at 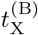 and died at 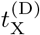, we exclude scenarios occurring in the range (*t*_1_, *t*_2_) that are beyond the lifetime of X, using a Heaviside step term

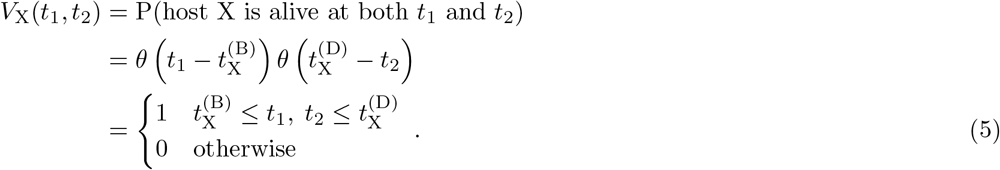

#### 4.3.4 Physical distance between hosts

Direct host-to-host transmission occurs from close contact, thus the relative transmission likelihood should depend on the separation between hosts (affecting how likely they are to interact). We capture this with a distance-dependence on the infection rate *β*. To account for the different movement patterns of wild badgers and domesticated cattle, we treat the interaction distance differently depending on the host species.

##### Cattle-cattle

For cattle-cattle transmission, we take the infection rate to be non-zero only when they were in the same herd at the same time. We identify interactions from their movement histories (CTS table `vMovementDirect`). The within-herd infection rate in those periods is taken to be that at zero separation. For all other times, the infection rate is zero:

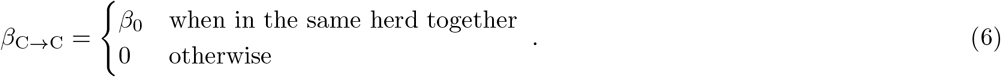

The relative likelihood of direct transmission between two cows that were never in the same herd at the same time is therefore zero. For *β*_0_ we use a gamma-distributed distribution based on the distribution obtained in Brooks-Pollock et al. [54], with a posterior estimate of the rate of within-holding onward transmission between cattle as *β*_0_ ∼ 0.61 y^−1^ (95% CI: [0.0503, 1.54]).

Also from cattle movement histories we identify *indirect* interactions between two cows. These are when the first cow interacted with one or more cows, that *later* went on to interact with the second, and could therefore have acted as an intermediary in a three-cow transmission chain.

For these interactions with intermediaries, the transmission rate is also taken to be that at zero distance. We multiply each relative likelihood by the number of intermediaries found in CTS, to estimate the relative likelihood of transmission via any one of these intermediaries.

##### Badger-badger

Medium-to-high density badger populations in the UK are typically organised into social groups with relatively well-defined territories [55, 56]. This was the case for Woodchester Park, where the social groups of individual badgers were closely monitored and group territories were well-defined during the period of study. The majority of movements and interactions between badgers have been observed to be within the territories of their own social group [57]. Between-social group movements are less frequent but serve important ecological purposes such as reducing inbreeding [58, 59]. The amount of between-group activity likely varies by badger depending on factors including age and sex [60, 61].

We take the effective rate of infection *β*_B→B_ between two badgers to fall exponentially with the separation *d* between them:

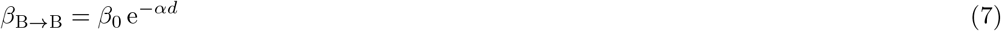

where *α* characterises how steep the fall with distance is. The social structure suggests that the majority of badger-badger transmission of *M. bovis* will be between badgers of the same social group, but it is more difficult to quantify the relative frequency of transmission pathways between badgers in adjacent social groups. Smith et al. [62] note that *M. bovis* infection originating specifically from a bite wound may be indicative of a between-group transmission, where the bite is inflicted by an infectious badger in a neighbouring social group, as a form of territorial aggression. From their study, of 33 infections 10 were from bite wounds, suggesting that up to 30% of infections may have been from inter-group pathways. If we estimate that a given social group will have approximately 5 social groups that could be considered adjacent, the proportion of infections within a social group from each neighbouring groups is 0.06.

Using the location data of the badgers at Woodchester Park, the mean distance between a social group and its 5 nearest neighbouring groups is 0.50 km. We then use a value *α* = 4.92 km^−1^ such that the infection rate at 0.50 km is 0.06/0.70 times that at zero distance.

##### Badger-cattle and cattle-badger

The cattle-badger and badger-cattle transmission rates also depend on distance, however the distance varies due to the movement of cattle between holdings:

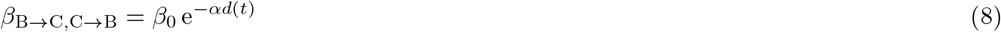

thus the infection rate changes in a step-like fashion whenever a cow moves holding, based on the physical distance between each holding and where the badger was observed. The same distance-dependency *α* is used as in Eq (7), reflecting the estimated roaming of the badgers. The physical locations of the holdings (used to determine a physical distance to the badger sampling location) are derived from CTS, table `tblLocation`.

#### 4.3.5 Evolutionary process of *M. bovis*

We assume that within-host single nucleotide polymorphisms (SNPs) of *M. bovis* are generated randomly and independently under Poisson dynamics, at a mean rate *μ*. Under Poisson dynamics, over an interval ∆*t*, the probability that *exactly m* SNPs are generated is

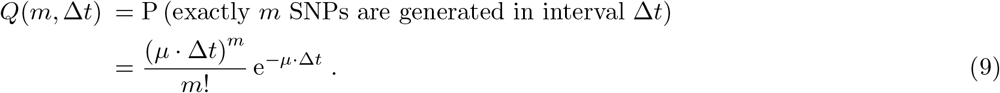

In an interval ∆*t* the expected number of SNPs to have been generated is ⟨ *m*⟩ = *μ*∆*t*, with standard deviation 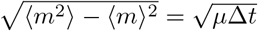. *M. bovis* exhibits a slow rate of molecular evolution; the substitution rate *μ* is estimated across different studies to be within the range of *μ*≈ [0.1 −0.9] substitutions, per genome, per year [11–15].

From the aggregation of all WGS samples, we identify the most common nucleotide per site, using this sequence of nucleotides as a reference. Then, for each pair of hosts A and B we count i) the number of divergent SNPs (differences between A’s sequence and the reference) observed in A but *not* in B (denoted *m*_A_), and; ii) the number of divergent SNPs observed in B but *not* in A (*m*_B_). Note that the sum *m*_A_ + *m*_B_ is the overall genetic distance between A and B, independent of the reference used. In a scenario where A infected B or vice versa, the infection event is the point of divergence for those two *M. bovis* strains, so any SNPs unique to one host only must have been generated *between* infection and observation (Fig 3).

Following Section 4.3.1, suppose the infection A → B occurred at time λ_B_. A and B are later sampled and their strains sequenced at times *t*_A_, *t*_B_ respectively. A’s sample has *m*_A_ SNPs not seen in B’s sample, and B’s has *m*_B_ SNPs not seen in A’s. By Eq (9), the probability of this is

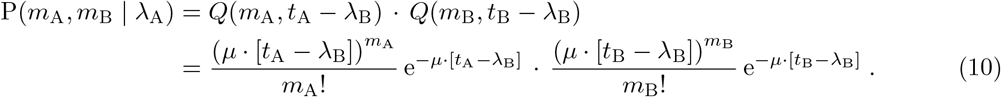

This incorporates the *different* sampling times of hosts, and the evolutionary processes that would happen *independently* in the two hosts post-infection.

Before incorporating WGS, the space of possible scenarios for transmission from one host to another is constrained by the host lifespans, and when those hosts interacted. The addition of WGS then constrains us to only scenarios where a specific number of SNPs are generated between infection and observation. We therefore expect the relative likelihoods for all pathways to fall compared to the equivalent calculation not including WGS data (which in effect sets the expression in Eq (9) to 1).

#### 4.3.6 Calculation

To evaluate the relative transmission likelihood between two hosts we numerically integrate over all possible times for the epidemiological events (Eqs (2), (3)), constrained by the delay to test sensitivity (Eq (4)) and the host lifespans (Eq (5)), with the infection rate dependent on the host distance (Eqs (6)–(8)). The relative likelihoods with WGS information are then further weighted by the probability of the observed genetic distance occurring (Eq (9)). Full expressions are presented in Supplementary Material, Section C.

For each transmission pathway, we evaluate the relative likelihood from 300 samples of the parameter distributions (Supplementary Material, Table S2). For the 250 host pairs with the highest median relative likelihood after this, we do a further 1000 evaluations, after which we record the median relative likelihood, and [0.05, 0.95] confidence interval.

Analysis code was written in *R* (version 4.3.1) [63]). To evaluate relative likelihoods we used multivariable numerical integration, with a nested application of the `integrate` function (`stats` package [63]). The implementation in *R* was verified against a symbolic implementation in *Mathematica* (version 12.3.1.0 [64]), using the `NIntegrate` function.

## 5. Acknowledgements

We thank Gianluigi Rossi for helpful discussions on relative transmission likelihoods, and providing comments on the manuscript.

## 6. Author contributions

RRK conceived the project. AJW performed formal analysis and with RRK wrote the manuscript. CMP and GM provided input on the statistical analysis, RJD and CHB on interpretation of the badger data, GCS on badger modelling, and EP on interpretation of the sequencing data. All authors commented on and approved the manuscript.

## 8. Code availability

Analysis code is available at https://git.ecdf.ed.ac.uk/awood310/mBovis-WGS-transmission-likelihoods.

## 9. Data availability

The cattle movement and testing data (CTS, SAM), the Woodchester Park study data and the WGS data are not publicly available, and are provided to the authors under a data sharing agreement with the Animal and Plant Health Agency, who can be contacted via.

## Supplementary Material

## A Spatial and temporal distribution of samples

**Figure S1:**
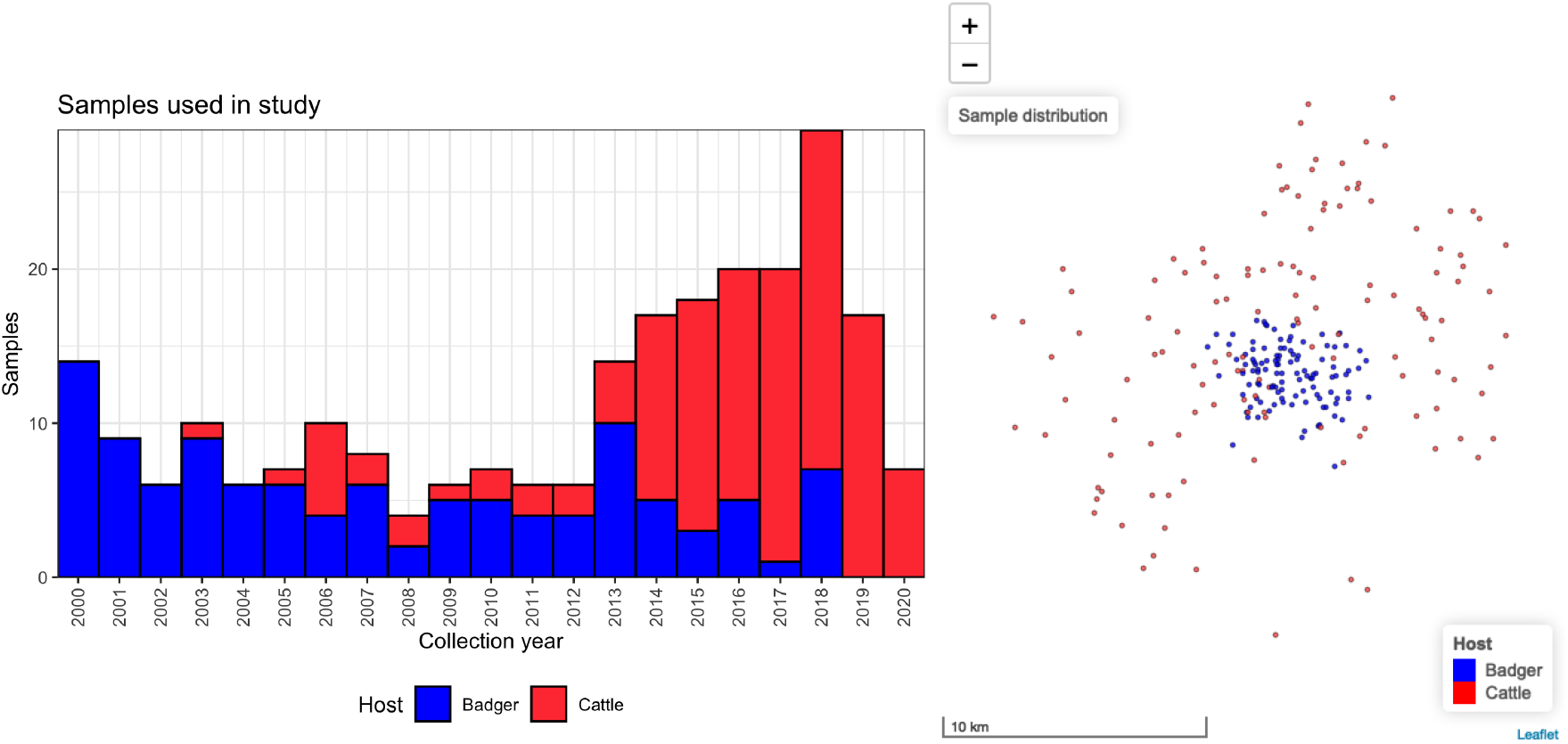
Distribution of samples used in this study. The locations (right) are of the hosts at the time the sequenced sample was taken. Badger samples (blue) are concentrated within the Woodchester Park study area, with cattle (red) in surrounding holdings. Exact locations have been jittered to reveal samples taken at the same location.

**Figure S2:**
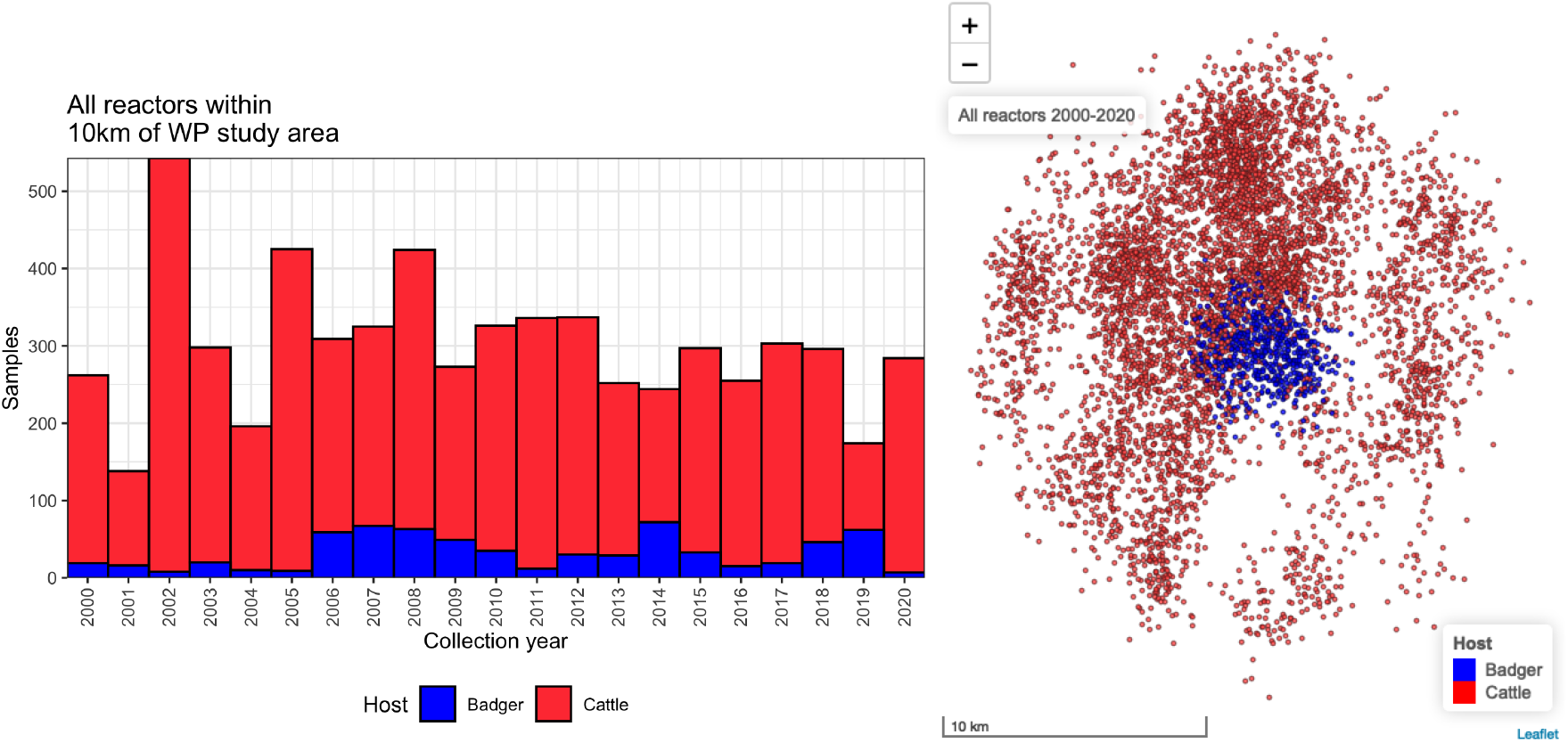
Distribution of *all* cattle reactors, and badgers with infection identified via culture or a serological test, within a 10 km radius of the WP study area between 2000–2020. Exact locations have been jittered to reveal samples taken at the same location.

## B Transmission pathways considered in this work

**Table S1:** List of potential transmission pathways where a first host is exposed, then goes on to expose a second host to infection either directly or indirectly, and which pathways are considered in this analysis. Note that any we only consider transmission pathways where the two hosts are members of the same direct lineage.

| Transmission pathway | Considered |
| --- | --- |
| Direct cow-cow transmission | Yes |
| Direct badger-badger transmission | Yes |
| Direct badger-cow transmission | Yes |
| Cow-cow transmission; chain with one intermediary cow | Yes |
| Cow-cow transmission; chain with one intermediary badger | No |
| Cow-badger transmission; chain with one intermediary host (any species) | No |
| Badger-badger transmission; chain with one intermediary host (any species) | No |
| Transmission chains involving >1 intermediary host, of any species | No |
| Transmission (any species) via an intermediary fomite | No |

## C Integral expressions for relative transmission likelihood

**Table S2:** Parameter ranges used in the relative likelihood calculation. The Γ(*p*_1_, *p*_2_) functions are gamma distributions with shape *p*_1_ and decay rate *p*_2_.

| Symbol | Description | Units | Distribution | Source |
| --- | --- | --- | --- | --- |
| $\beta_0$ | Infection rate within herd/at zero distance | $y^{-1}$ | $\Gamma(1.60, 2.63)$ | [54] |
| $\alpha$ | Exponential rate of decay of infection rate with distance | $\text{km}^{-1}$ | 4.92 | [62] |
| $\sigma$ | Rate of transition to infectiousness | $y^{-1}$ | $\Gamma(4.44, 0.40)$ | [54] |
| $\Omega$ | Time between infection and test-sensitivity | $y$ | $U(0, 0.23)$ | [53] |
| $\mu$ | Rate of *M. bovis* substitution events per genome | $y^{-1}$ | $U(0.1, 0.9)$ | [11–14] |

In this section we describe a method for evaluating the relative likelihood that a transmission occurred from one host to another. Recalling Section 4.3 this is informed by:

- the *epidemiology* of bTB and the timescales of the different epidemiological processes;
- the time between exposure, and detectability by diagnostic test;
- the known *lifespans* of the badgers and cattle;
- the known *locations* of the badgers and cattle;
- the *evolutionary* characteristics of *M. bovis*.

We label the two hosts of interest A and B, choosing A as the host sampled first. We consider scenarios where:

- A directly infected B (ℒ_A→B_, Fig 3A);
- B directly infected A (ℒ_B→A_, Fig 3B);
- (cattle-cattle only) A infected an intermediary X, that later infected B (ℒ_A→X→B_, Fig S3A)
- (cattle-cattle only) B infected an intermediary X, that later infected A (ℒ_B→X→A_, Fig S3B).

### C.1 Relative transmission likelihoods informed by life history data and WGS

#### ℒ_*A*→*B*_**; A exposes B directly**

We observe that A and B were test-sensitive at times *t*_A_, *t*_B_, where A had *m*_A_ unique SNPs at *t*_A_ not seen in B’s WGS sample; and B had *m*_B_ unique SNPs not seen in A’s WGS sample.

For A to have directly exposed B, the following chain of epidemiological events must have occurred:

1. A was exposed at some time λ_A_ (by an unknown host);
2. A became infectious at some time *ξ*_A_;
3. A exposed B at some time λ_B_.

This chain forces λ_A_ ≤*ξ*_A_ ≤λ_B_. These are further constrained by the observation times *t*_A_, *t*_B_, the delay to test-sensitivity (Eq (4)), and host lifespans (Eq (5)).

The WGS observation then requires that A generated *m*_A_ SNPs between infecting B and being observed at *t*_A_, and that B generated *m*_B_ SNPs between being exposed by A, and being observed at *t*_B_.

Summarising,

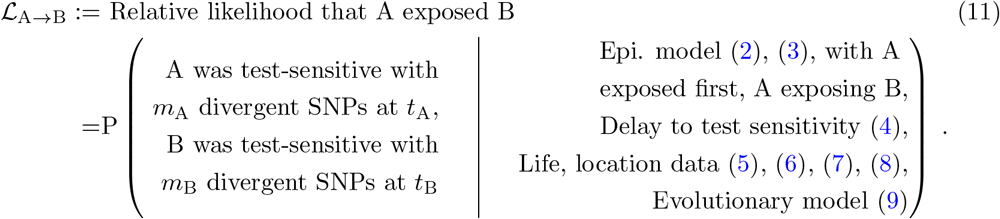

We integrate over all possible times λ_A_, *ξ*_A_, λ_B_ in the epidemiological chain, as weighted by the conditional probability densities in Eqs (2), (3). We exclude scenarios incompatible with the lifespans and delay to test sensitivity with the terms (Eqs (4), (5)).

Finally we weight scenarios with the probability of the observed number of divergent SNPs being generated, Eq (9). Care is required to differentiate between SNPs that would only be unique to one host, and SNPs what would be generated by A but then inherited by B, thus being shared. A can generate SNPs from λ_A_. However, SNPs generated *prior* to λ_B_ (the time that A exposes B) are shared by both, thus not divergent. This gives three possible scenarios:

1. A exposes B prior to *t*_A_. Both A and B can generate divergent SNPs after λ_B_. The relative likelihood of this scenario is:

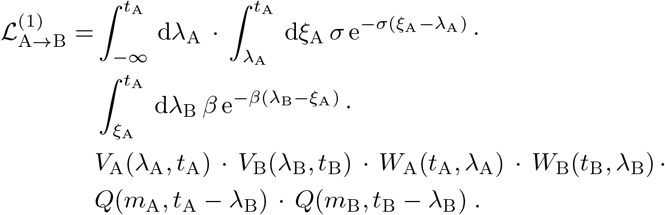 Note the lower bound for the λ_A_ integral is −∞ (saying A could have been first exposed at any point before being observed), though this is trivially bound by the lifespan of A *V*_A_(λ_A_, *t*_A_) (5). As we integrate over a range of possible times for A’s initial exposure without weighting, it is possible for the relative likelihoods as defined in this work to exceed 1.
2. A became infectious before *t*_A_, but did not expose B until after *t*_A_ (this is mainly applicable to badgers, as cattle are slaughtered shortly after a positive test). A could not generate divergent SNPs before observation, B can only generate divergent SNPs past λ_B_, or inherit SNPs generated by A (but not observed) between *t*_A_ and λ_B_:

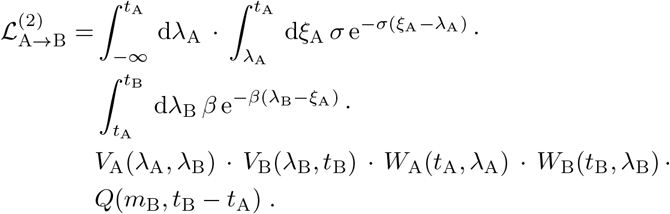
3. A became infectious after *t*_A_, and B was not exposed until after *t*_A_. A could not generate divergent SNPs before observation, B can only generate divergent SNPs past λ_B_, or inherit SNPs generated by A (but not observed) between *t*_A_ and λ_B_:

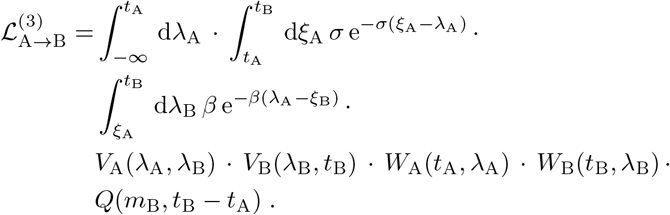

The full relative likelihood ℒ_A→B_ is a summation of all three terms:

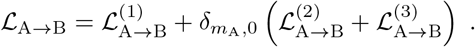

#### ℒ_B→A_, **B exposes A directly**

Suppose now that the infection occurred in the other direction; B was first exposed at λ_B_, became infectious at *ξ*_B_ and exposed A at λ_A_. The observations at *t*_A_ and *t*_B_ bound the time λ_A_ that A could have been exposed by B: λ_B_ ≤ λ_A_ ≤ *t*_A_. The relative likelihood is then

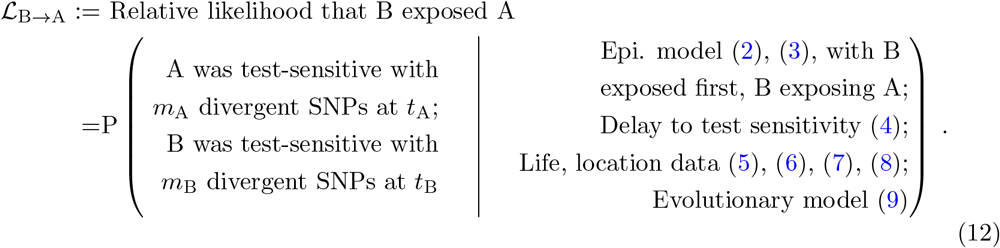

This is identical to Eq (11) except B is exposed first in the epidemiological model.

As A was observed before B, the exposure must have happened before *t*_A_. The relative likelihood that B exposed A is then a single term:

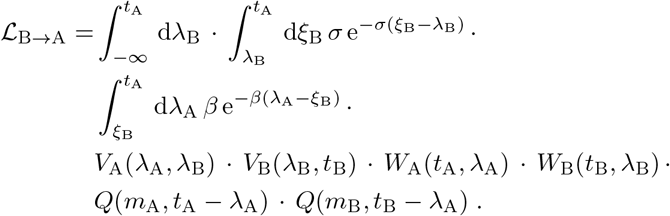

#### ℒ_A→X→B_**; A exposes B via an intermediary X (cattle-to-cattle only)**

For cattle-cattle transmission, we also consider the presence of an intermediary cow X. The chain of events is then:

**Figure S3:**
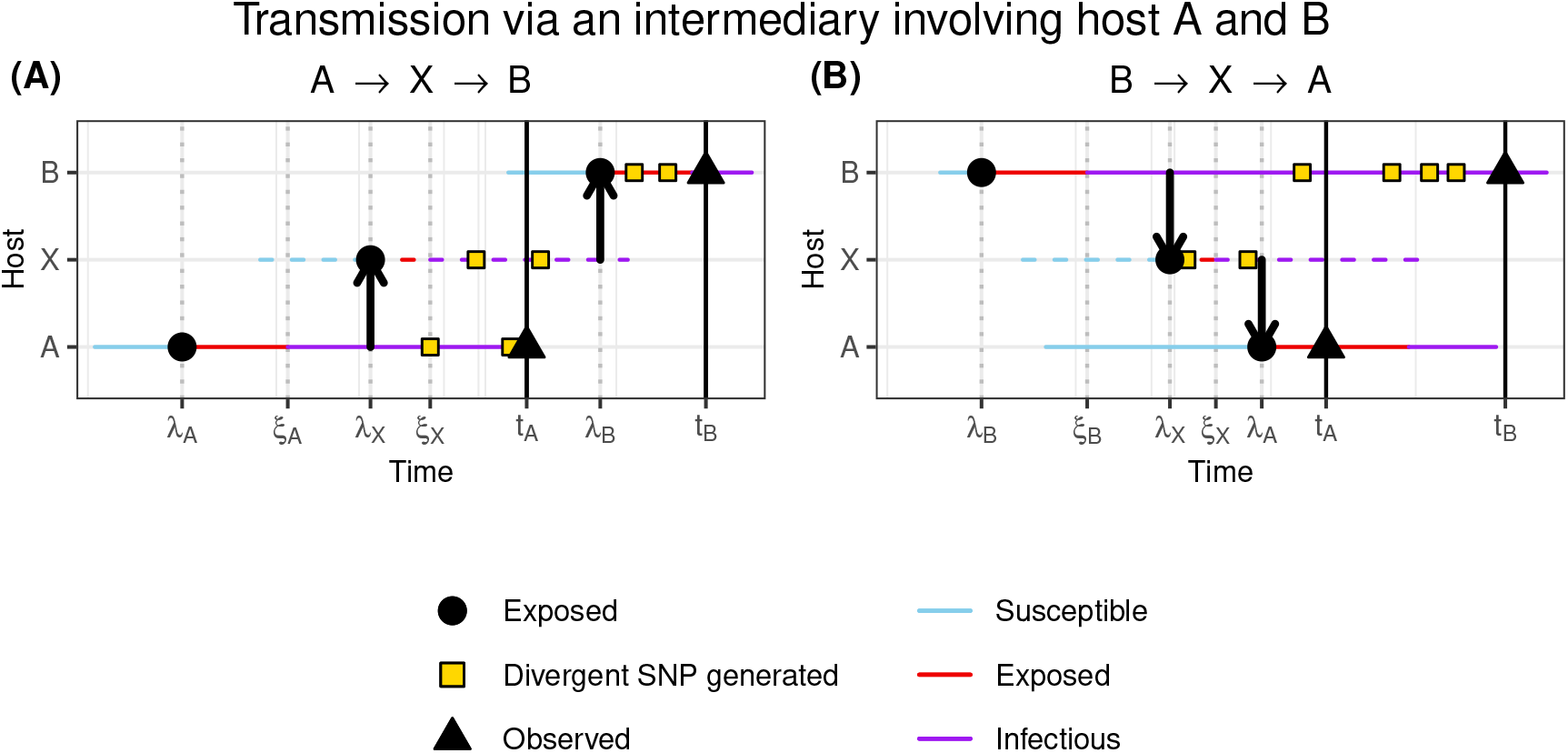
For two cattle hosts A and B that are sampled at times *t*_A_ and *t*_B_ respectively, two transmission pathways via an unobserved intermediary cow X that would be consistent with an observed genetic distance of 6 SNPs, with A having 2 SNPs not seen in B, and B with 4 SNPs not seen in A, both with respect to a reference genome. Hosts are born susceptible (blue), exposed to infection at times denoted λ (red), and become infectious at *ξ* (purple). The generation of SNPs on the pathogen genome inside each host are denoted with yellow squares. (**A**): A exposes X to infection, who goes on to expose B to infection. (**B**): B exposes X to infection, who goes on to expose A to infection.

1. A was exposed at some time λ_A_ (by an unknown host);
2. A became infectious at some time *ξ*_A_;
3. A exposed X at some time λ_X_;
4. X became infectious at some time *ξ*_X_;
5. X exposed B at some time λ_B_.

This chain forces λ_A_ ≤ *ξ*_A_ ≤ λ_X_ ≤ *ξ*_X_ ≤ λ_B_. Given the same observations at *t*_A_ and *t*_B_, the relative likelihood is a generalisation of the A→B system:

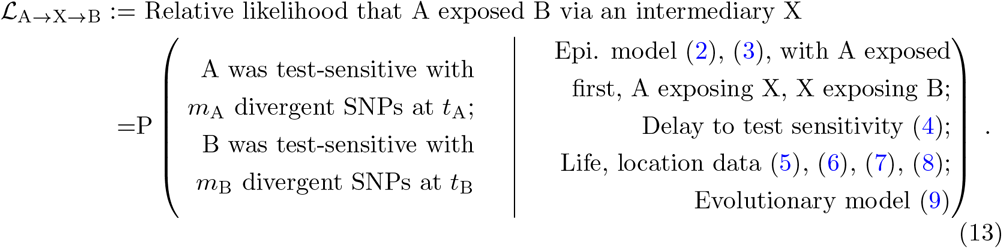

As with ℒ _A→B_ we need to take care to track whether SNPs generated by the hosts are unique to one host, or inherited by the second and therefore not divergent. We again have three scenarios:

1. A exposes X prior to *t*_A_. A and X can generate divergent SNPs after λ_X_. B inherits SNPs generated by X between λ_X_ and λ_B_, and can also generate SNPs between λ_B_ and *t*_B_:

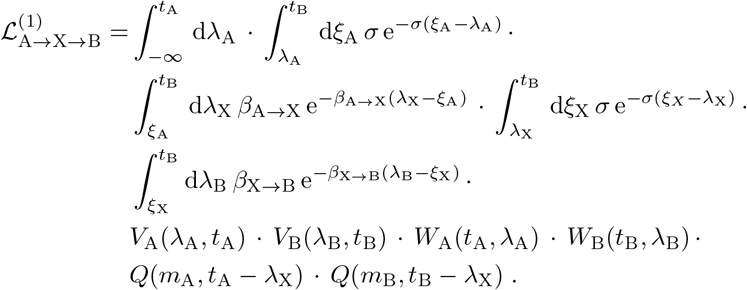

We have separated the infection rate *β*_A→X_ from host A to X, and the infection rate *β*_X→B_ from host X to B.

2. A became infectious before *t*_A_, but did not expose X until after. A could not generate divergent SNPs before observation, X can only generate divergent SNPs past λ_X_, or inherit SNPs generated by X between *t*_A_ and λ_X_. B can generate SNPs past λ_B_, and inherit any from X:

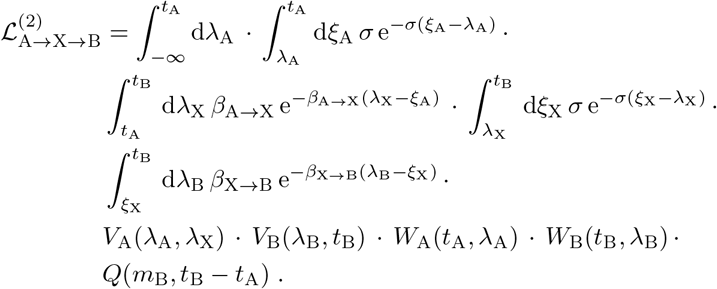

3. A only became infectious after *t*_A_, thus X was not exposed until after *t*_A_. A could not generate divergent SNPs before observation, X can only generate divergent SNPs past λ_X_, or inherit SNPs generated by A (but not observed) between *t*_A_ and λ_X_. B can generate SNPs past λ_B_, and inherit any from X:

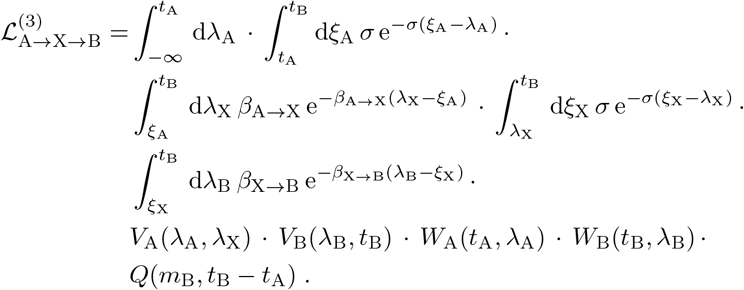

The relative likelihood ℒ_A→X→B_ is then a summation of these terms:

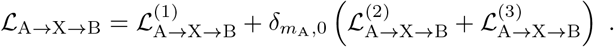

#### ℒ_B→X→A_**; B exposes A via an intermediary X (cattle-to-cattle only)**

Finally, suppose that B was first exposed at λ_B_, became infectious at *ξ*_B_ and exposed X at time λ_X_, who later exposes A at time λ_A_:

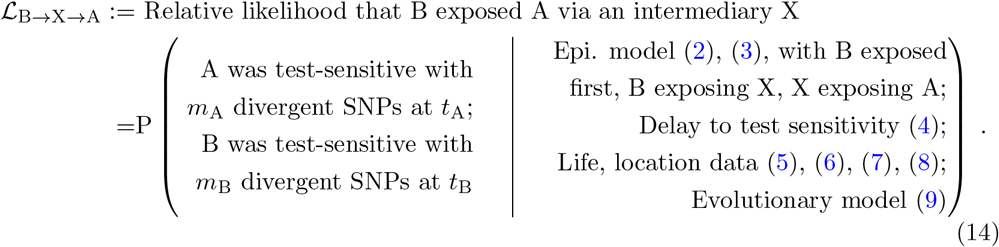

Given the same observations, this bounds the time that A could have been exposed by X: λ_B_ ≤ λ_A_ ≤ *t*_A_. The relative likelihood is then:

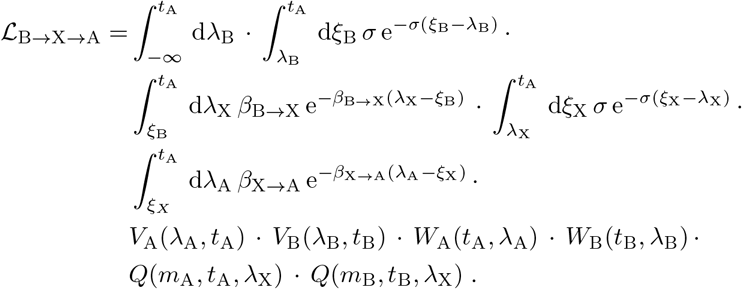

### Relative transmission likelihoods on the basis of life history data alone

The expressions for relative transmission likelihoods *not* informed by WGS and only on the basis of life history alone are equivalent to those presented, with the evolutionary terms *Q* are set to 1.

## D Relative transmission likelihoods; per-species, overall distribution

**Figure S4:**
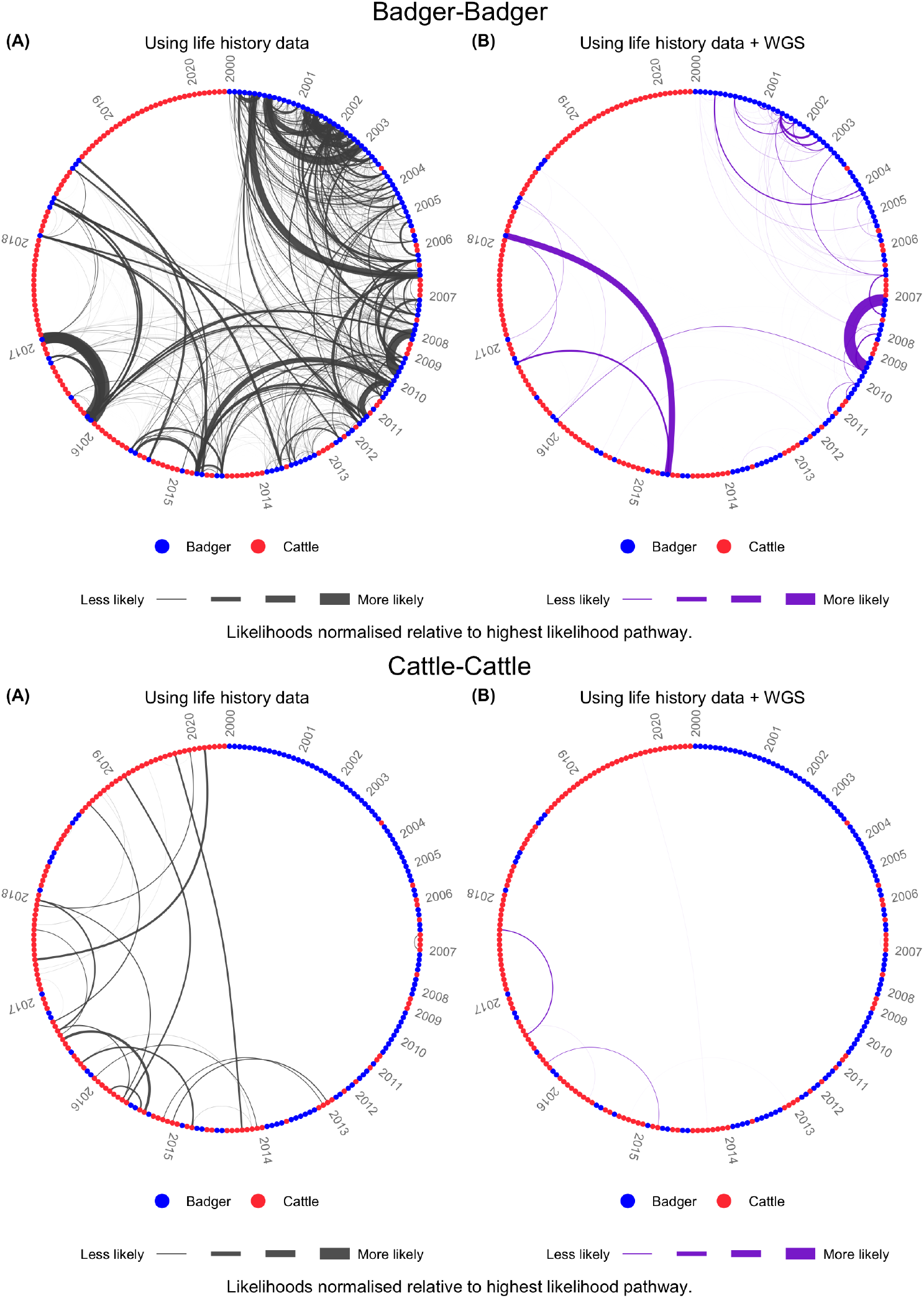
Relative transmission likelihoods from Fig 1, isolating badger-badger (top) and cattlecattle (bottom) transmission pathways only.

**Figure S5:**
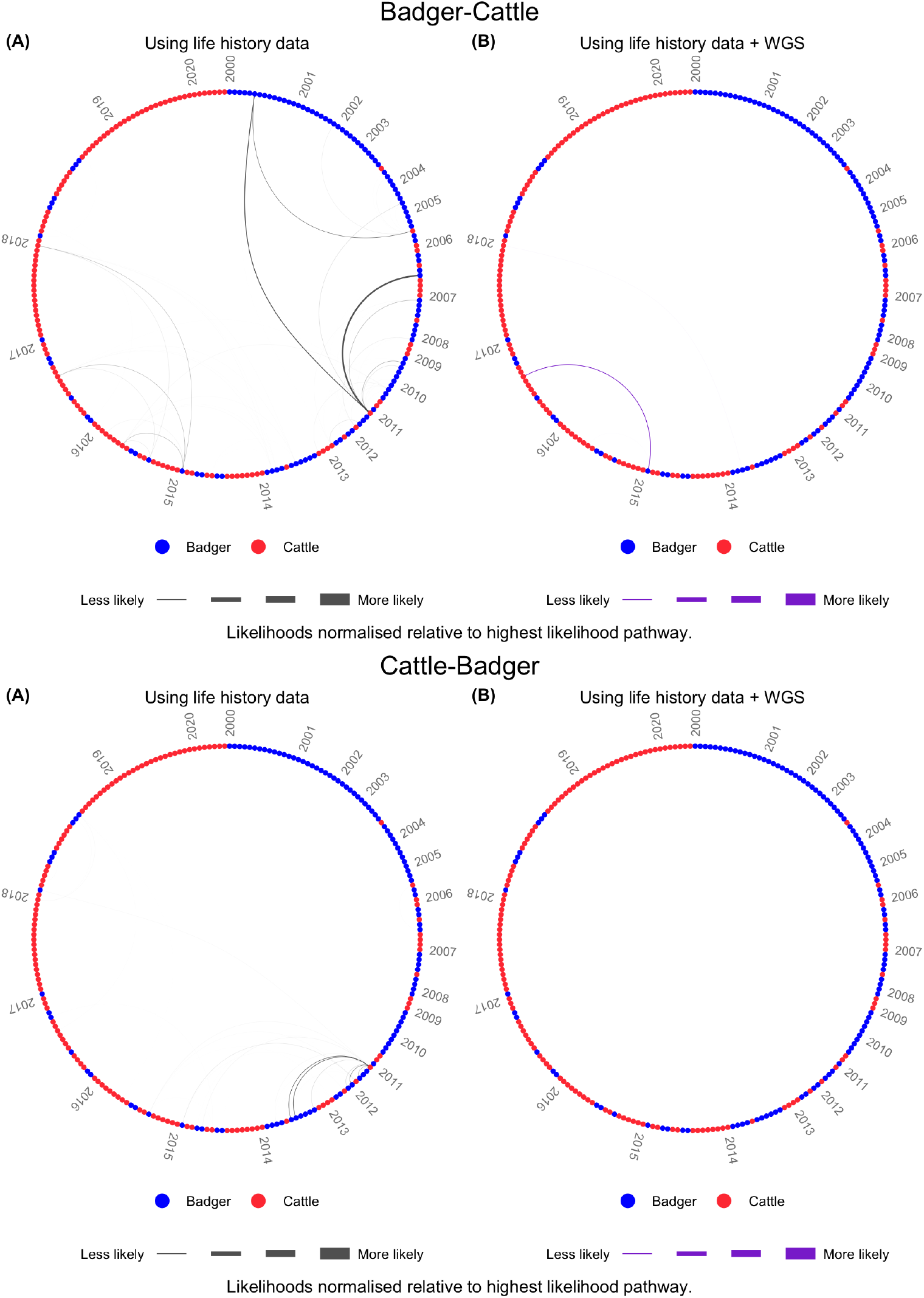
Relative transmission likelihoods from Fig 1, isolating badger-cattle (top) and cattle-badger (bottom) transmission pathways only.

**Figure S6:**
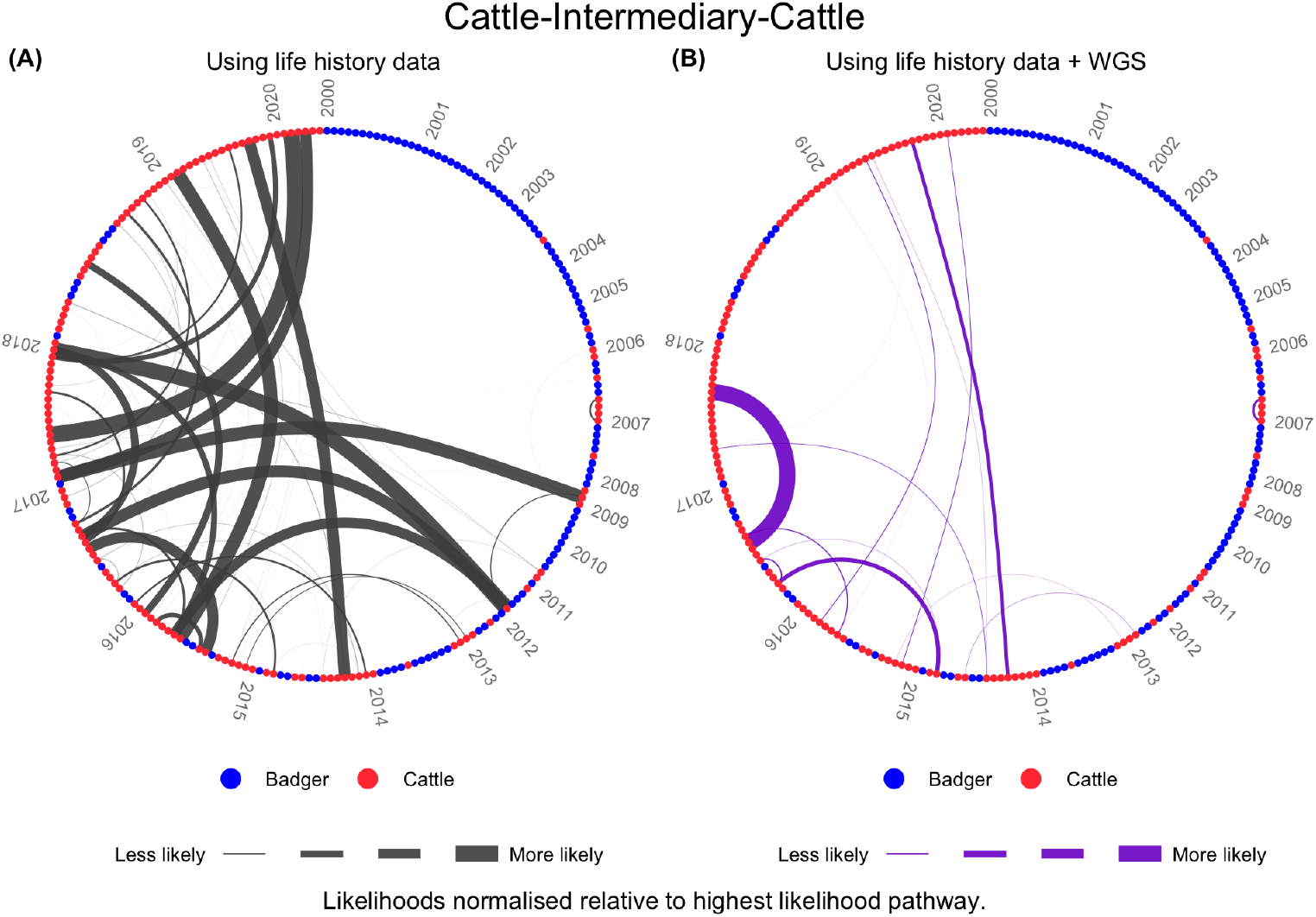
Relative transmission likelihoods over pathways between cattle pairs including infection via an intermediary cattle host.

**Figure S7:**
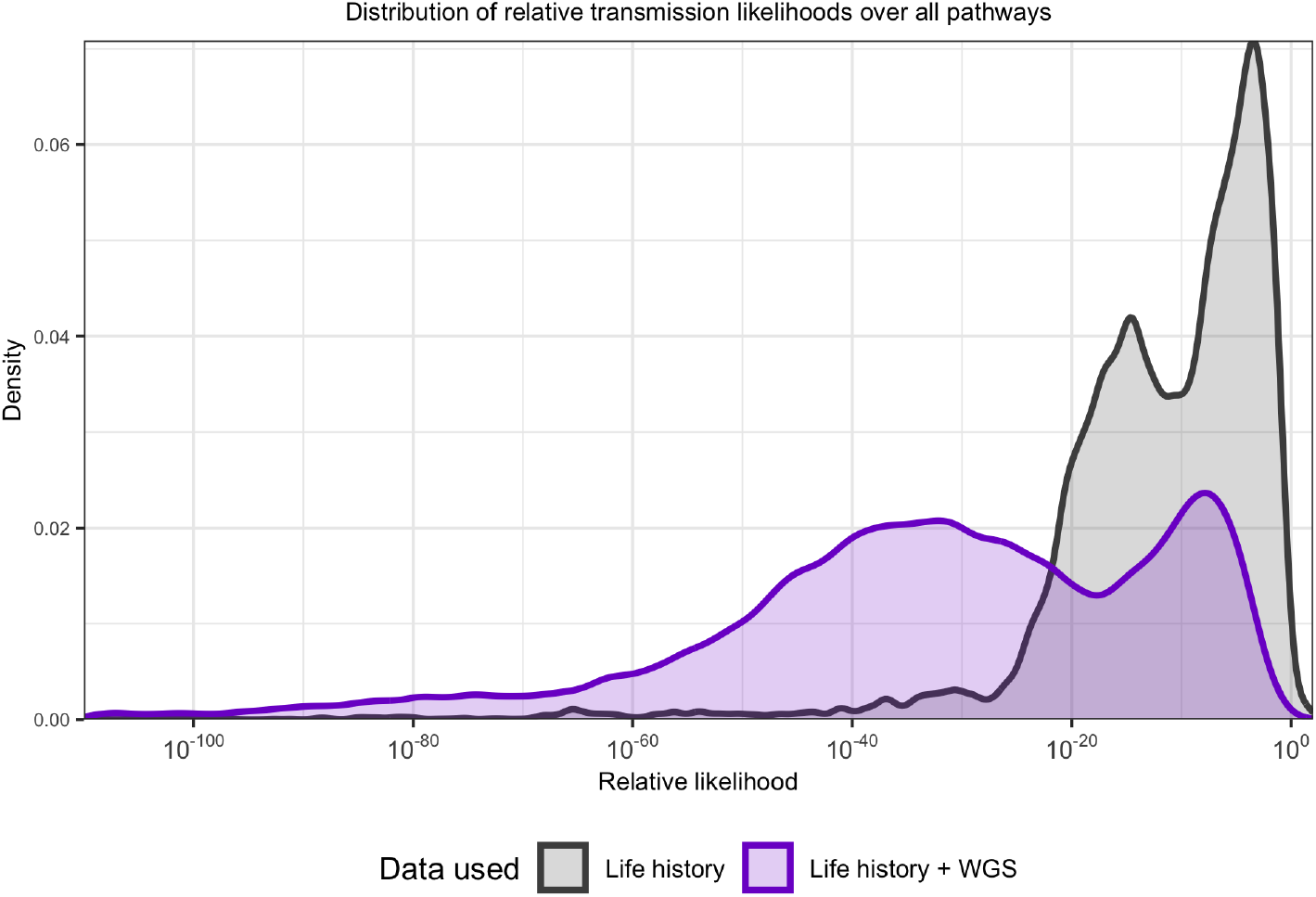
The distribution of relative likelihoods of all pathways from Figure 1 (noting log scale on the *x*-axis)

## E Relative transmission likelihood for specific host pairs

**Table S3:** Transmission pathways fitting category *(i)* in Table 1, having a high relative likelihood on the basis of life history data, and corroborated by the additional WGS data.

| Pathway | Lifespan<br>(infector) | Lifespan<br>(infected) | Sample<br>time<br>(infector) | Sample<br>time<br>(infected) | Physical<br>distance<br>(km) | Relative<br>likelihood<br>(LH) | Genetic<br>distance<br>(SNPs) | Relative<br>likelihood<br>(LH + WGS) |
| --- | --- | --- | --- | --- | --- | --- | --- | --- |
| C.028>X>C.120 | 2009.2–2014.2 | 2011.6–2019.9 | 2014.2 | 2019.9 | - | 36.6823 (rank 6) | 5 | 0.13659 (rank 5) |
| B.019>B.052 | 1993.1–2002.2 | 1999.1–2007.1 | 2001.5 | 2006.4 | 0.4 | 0.3732 (rank 91) | 8 | 0.00096 (rank 94) |
| B.069>B.074 | 2004.1–2011.4 | 2007.1–2012.0 | 2010.4 | 2011.4 | 0.0 | 0.9694 (rank 51) | 5 | 0.00505 (rank 47) |
| C.061>X>C.045 | 2012.8–2016.8 | 2012.5–2015.8 | 2016.7 | 2015.8 | - | 1.6187 (rank 39) | 5 | 0.00988 (rank 34) |
| C.045>X>C.061 | 2012.5–2015.8 | 2012.8–2016.8 | 2015.8 | 2016.7 | - | 7.6600 (rank 19) | 5 | 0.03882 (rank 12) |
| C.039>X>C.021 | 2006.0–2015.3 | 2009.4–2013.2 | 2015.2 | 2013.2 | - | 10.0866 (rank 17) | 7 | 0.12393 (rank 7) |
| C.054>X>C.034 | 2009.1–2016.3 | 2011.0–2015.0 | 2016.2 | 2014.9 | - | 11.4278 (rank 16) | 4 | 0.40166 (rank 2) |
| C.021>X>C.039 | 2009.4–2013.2 | 2006.0–2015.3 | 2013.2 | 2015.2 | - | 3.7204 (rank 29) | 7 | 0.00673 (rank 44) |
| B.061>B.066 | 1999.1–2008.5 | 2004.1–2010.3 | 2008.0 | 2009.8 | 0.0 | 1.5481 (rank 41) | 4 | 0.01326 (rank 25) |
| B.009>B.022 | 1993.1–2002.2 | 1998.1–2002.6 | 2000.5 | 2001.6 | 0.1 | 0.9147 (rank 52) | 4 | 0.00921 (rank 36) |
| C.039>C.021 | 2006.0–2015.3 | 2009.4–2013.2 | 2015.2 | 2013.2 | - | 0.4378 (rank 84) | 7 | 0.00211 (rank 68) |
| C.034>X>C.054 | 2011.0–2015.0 | 2009.1–2016.3 | 2014.9 | 2016.2 | - | 6.9041 (rank 21) | 4 | 0.16366 (rank 4) |
| C.060>X>C.079 | 2013.8–2016.7 | 2013.7–2017.9 | 2016.6 | 2017.8 | - | 7.1052 (rank 20) | 2 | 0.62861 (rank 1) |
| C.006>X>C.009 | 2001.1–2006.6 | 2004.2–2007.3 | 2006.6 | 2007.3 | - | 5.4018 (rank 27) | 5 | 0.08984 (rank 8) |
| C.028>C.120 | 2009.2–2014.2 | 2011.6–2019.9 | 2014.2 | 2019.9 | - | 0.2833 (rank 121) | 5 | 0.00036 (rank 141) |

**Table S4:** Transmission pathways fitting category *(ii)* in Table 1, having a very low relative likelihood on the basis of life history data, and corroborated by the additional WGS data.

| Pathway | Lifespan<br>(infector) | Lifespan<br>(infected) | Sample<br>time<br>(infector) | Sample<br>time<br>(infected) | Physical<br>distance<br>(km) | Relative<br>likelihood<br>(LH) | Genetic<br>distance<br>(SNPs) | Relative<br>likelihood<br>(LH + WGS) |
| --- | --- | --- | --- | --- | --- | --- | --- | --- |
| B.090>C.023 | 2013.1–2014.9 | 2012.3–2013.6 | 2013.8 | 2013.6 | 8.2 | 1.0e-21 (rank 10675) | 49 | 1.8e-101 (rank 10676) |
| B.108>C.112 | 2016.1–2019.1 | 2017.8–2019.6 | 2018.5 | 2019.6 | 9.2 | 1.1e-23 (rank 11023) | 30 | 8.8e-202 (rank 11020) |
| B.111>C.089 | 2015.1–2019.9 | 2016.6–2018.3 | 2018.7 | 2018.2 | 10.1 | 4.5e-23 (rank 10932) | 91 | 1.1e-167 (rank 10937) |
| B.108>C.119 | 2016.1–2019.1 | 2018.6–2019.9 | 2018.5 | 2019.9 | 9.3 | 8.6e-22 (rank 10693) | 40 | 9.7e-106 (rank 10702) |
| C.089>B.106 | 2016.6–2018.3 | 2018.1–2020.0 | 2018.2 | 2018.4 | 9.8 | 1.2e-23 (rank 11020) | 88 | 8.9e-215 (rank 11040) |
| B.096>C.076 | 2007.1–2016.0 | 2015.8–2017.7 | 2015.5 | 2017.7 | 9.6 | 4.0e-22 (rank 10773) | 31 | 3.2e-131 (rank 10807) |
| B.102>C.089 | 2013.1–2017.6 | 2016.6–2018.3 | 2016.8 | 2018.2 | 10.1 | 5.3e-23 (rank 10920) | 85 | 1.5e-174 (rank 10954) |
| B.083>C.127 | 2013.1–2015.1 | 2013.4–2020.2 | 2013.5 | 2020.2 | 10.2 | 9.6e-25 (rank 11157) | 31 | 2.5e-280 (rank 11206) |
| B.090>C.104 | 2013.1–2014.9 | 2008.4–2019.0 | 2013.8 | 2018.9 | 9.0 | 2.1e-21 (rank 10588) | 48 | 6.8e-88 (rank 10532) |
| C.089>B.108 | 2016.6–2018.3 | 2016.1–2019.1 | 2018.2 | 2018.5 | 9.6 | 8.7e-23 (rank 10889) | 93 | 1.4e-171 (rank 10946) |
| B.089>C.038 | 2012.1–2014.2 | 2013.1–2015.2 | 2013.7 | 2015.2 | 7.2 | 1.2e-27 (rank 11325) | 12 | 1.9e-295 (rank 11264) |
| B.044>B.036 | 2004.1–2007.1 | 2002.1–2004.9 | 2004.7 | 2003.9 | 0.8 | 7.7e-24 (rank 11044) | 113 | 7.5e-241 (rank 11108) |
| B.044>B.037 | 2004.1–2007.1 | 2003.1–2004.9 | 2004.7 | 2003.9 | 1.3 | 6.0e-25 (rank 11182) | 113 | 2.3e-242 (rank 11114) |
| B.107>C.119 | 2010.1–2019.0 | 2018.6–2019.9 | 2018.5 | 2019.9 | 9.8 | 6.0e-22 (rank 10730) | 38 | 1.5e-99 (rank 10660) |
| B.084>C.127 | 2013.1–2015.2 | 2013.4–2020.2 | 2013.5 | 2020.2 | 10.2 | 2.0e-24 (rank 11128) | 31 | 2.5e-280 (rank 11205) |

**Table S5:** Transmission pathways fitting category *(iii)* in Table 1, having a high relative likeli ood on the basis of life history data, but contradicted by the additional WGS data; the inclusion of WGS data decreases the relative likelihood compared to other pathways.

| Pathway | Lifespan<br>(infector) | Lifespan<br>(infectee) | Sample<br>time<br>(infector) | Sample<br>time<br>(infected) | Physical<br>distance<br>(km) | Relative<br>likelihood<br>(LH) | Genetic<br>distance<br>(SNPs) | Relative<br>likelihood<br>(LH + WGS) |
| --- | --- | --- | --- | --- | --- | --- | --- | --- |
| B.043>B.058 | 2003.1–2010.9 | 2004.1–2007.9 | 2004.7 | 2007.4 | 0.0 | 2.0e-01 (rank 171) | 149 | 2.9e-306 (rank 11288) |
| B.043>B.069 | 2003.1–2010.9 | 2004.1–2011.4 | 2004.7 | 2010.4 | 0.7 | 4.5e-02 (rank 454) | 149 | 1.5e-307 (rank 11294) |
| B.054>B.043 | 1997.1–2009.3 | 2003.1–2010.9 | 2006.6 | 2004.7 | 0.5 | 7.8e-02 (rank 327) | 149 | 2.2e-260 (rank 11154) |
| B.033>B.043 | 2001.1–2004.1 | 2003.1–2010.9 | 2003.4 | 2004.7 | 0.3 | 3.6e-02 (rank 508) | 157 | 9.3e-286 (rank 11225) |
| B.044>B.069 | 2004.1–2007.1 | 2004.1–2011.4 | 2004.7 | 2010.4 | 0.0 | 5.3e-02 (rank 403) | 113 | 2.7e-231 (rank 11063) |
| B.036>B.043 | 2002.1–2004.9 | 2003.1–2010.9 | 2003.9 | 2004.7 | 0.3 | 2.8e-02 (rank 571) | 149 | 4.9e-268 (rank 11167) |
| B.052>B.044 | 1999.1–2007.1 | 2004.1–2007.1 | 2006.4 | 2004.7 | 0.4 | 4.3e-02 (rank 466) | 117 | 2.1e-228 (rank 11056) |
| B.043>B.054 | 2003.1–2010.9 | 1997.1–2009.3 | 2004.7 | 2006.6 | 0.5 | 1.5e-02 (rank 779) | 149 | 3.3e-265 (rank 11163) |
| B.041>B.043 | 2000.1–2005.0 | 2003.1–2010.9 | 2004.5 | 2004.7 | 0.8 | 1.2e-02 (rank 859) | 149 | 1.4e-266 (rank 11166) |
| B.038>B.043 | 1998.1–2004.4 | 2003.1–2010.9 | 2003.9 | 2004.7 | 0.9 | 9.6e-03 (rank 925) | 149 | 7.9e-269 (rank 11170) |
| B.043>B.050 | 2003.1–2010.9 | 2004.1–2006.0 | 2004.7 | 2005.8 | 0.6 | 6.3e-03 (rank 1059) | 147 | 1.4e-309 (rank 11300) |
| B.043>B.051 | 2003.1–2010.9 | 2004.1–2006.8 | 2004.7 | 2006.4 | 0.8 | 3.6e-03 (rank 1219) | 151 | 5.5e-313 (rank 11308) |
| B.052>B.043 | 1999.1–2007.1 | 2003.1–2010.9 | 2006.4 | 2004.7 | 1.0 | 5.2e-03 (rank 1106) | 151 | 1.9e-265 (rank 11164) |
| B.039>B.043 | 2003.1–2004.5 | 2003.1–2010.9 | 2004.1 | 2004.7 | 0.3 | 4.9e-03 (rank 1131) | 149 | 8.6e-273 (rank 11186) |
| B.044>B.052 | 2004.1–2007.1 | 1999.1–2007.1 | 2004.7 | 2006.4 | 0.4 | 7.0e-03 (rank 1020) | 117 | 4.7e-233 (rank 11071) |

**Table S6:** Transmission pathways fitting category *(iv)* in Table 1, having a very low relative likelihood on the basis of life history data, but contradicted by the additional WGS data; the inclusion of WGS data increases the relative likelihood compared to other pathways.

| Pathway | Lifespan<br>(infector) | Lifespan<br>(infectee) | Sample<br>time<br>(infector) | Sample<br>time<br>(infected) | Physical<br>distance<br>(km) | Relative<br>likelihood<br>(LH) | Genetic<br>distance<br>(SNPs) | Relative<br>likelihood<br>(LH + WGS) |
| --- | --- | --- | --- | --- | --- | --- | --- | --- |
| B.102>C.056 | 2013.1–2017.6 | 2012.4–2016.4 | 2016.8 | 2016.3 | 5.2 | 1.8e-12 (rank 6827) | 3 | 1.9e-14 (rank 2403) |
| B.077>C.077 | 2010.1–2012.9 | 2008.5–2017.8 | 2012.4 | 2017.7 | 4.4 | 1.6e-11 (rank 6448) | 2 | 2.4e-12 (rank 2028) |
| B.012>B.002 | 2000.1–2001.3 | 1995.1–2000.6 | 2000.7 | 2000.1 | 0.3 | 2.1e-12 (rank 6804) | 2 | 1.5e-14 (rank 2428) |
| C.056>B.102 | 2012.4–2016.4 | 2013.1–2017.6 | 2016.3 | 2016.8 | 5.2 | 3.4e-12 (rank 6721) | 3 | 3.2e-14 (rank 2355) |
| B.097>C.056 | 2014.1–2016.4 | 2012.4–2016.4 | 2015.6 | 2016.3 | 3.9 | 2.7e-10 (rank 5955) | 0 | 1.7e-10 (rank 1622) |
| C.077>B.077 | 2008.5–2017.8 | 2010.1–2012.9 | 2017.7 | 2012.4 | 4.4 | 3.6e-11 (rank 6305) | 2 | 3.8e-12 (rank 1995) |
| B.108>C.129 | 2016.1–2019.1 | 2018.2–2020.4 | 2018.5 | 2020.4 | 2.5 | 7.0e-12 (rank 6596) | 4 | 7.2e-14 (rank 2303) |
| C.056>B.097 | 2012.4–2016.4 | 2014.1–2016.4 | 2016.3 | 2015.6 | 3.9 | 4.2e-10 (rank 5875) | 0 | 2.0e-10 (rank 1605) |
| B.094>C.048 | 2008.1–2015.9 | 2013.3–2015.9 | 2014.8 | 2015.9 | 5.5 | 4.7e-12 (rank 6668) | 4 | 1.6e-14 (rank 2414) |
| B.109>C.056 | 2015.1–2019.2 | 2012.4–2016.4 | 2018.7 | 2016.3 | 4.0 | 3.4e-11 (rank 6312) | 2 | 1.4e-12 (rank 2076) |
| B.094>C.105 | 2008.1–2015.9 | 2012.9–2019.0 | 2014.8 | 2018.9 | 5.5 | 4.3e-12 (rank 6677) | 7 | 1.2e-14 (rank 2450) |
| B.066>C.073 | 2004.1–2010.3 | 2008.6–2017.6 | 2009.8 | 2017.6 | 5.4 | 3.3e-12 (rank 6723) | 9 | 3.4e-15 (rank 2538) |
| B.076>C.077 | 2005.1–2011.9 | 2008.5–2017.8 | 2011.4 | 2017.7 | 4.4 | 4.3e-12 (rank 6676) | 9 | 6.5e-15 (rank 2498) |
| B.039>B.031 | 2003.1–2004.5 | 2000.1–2003.9 | 2004.1 | 2003.4 | 1.9 | 2.7e-09 (rank 5530) | 0 | 1.8e-09 (rank 1371) |
| C.019>B.066 | 2009.3–2012.7 | 2004.1–2010.3 | 2012.7 | 2009.8 | 3.5 | 1.4e-11 (rank 6473) | 6 | 2.0e-14 (rank 2395) |

**Figure S8:**
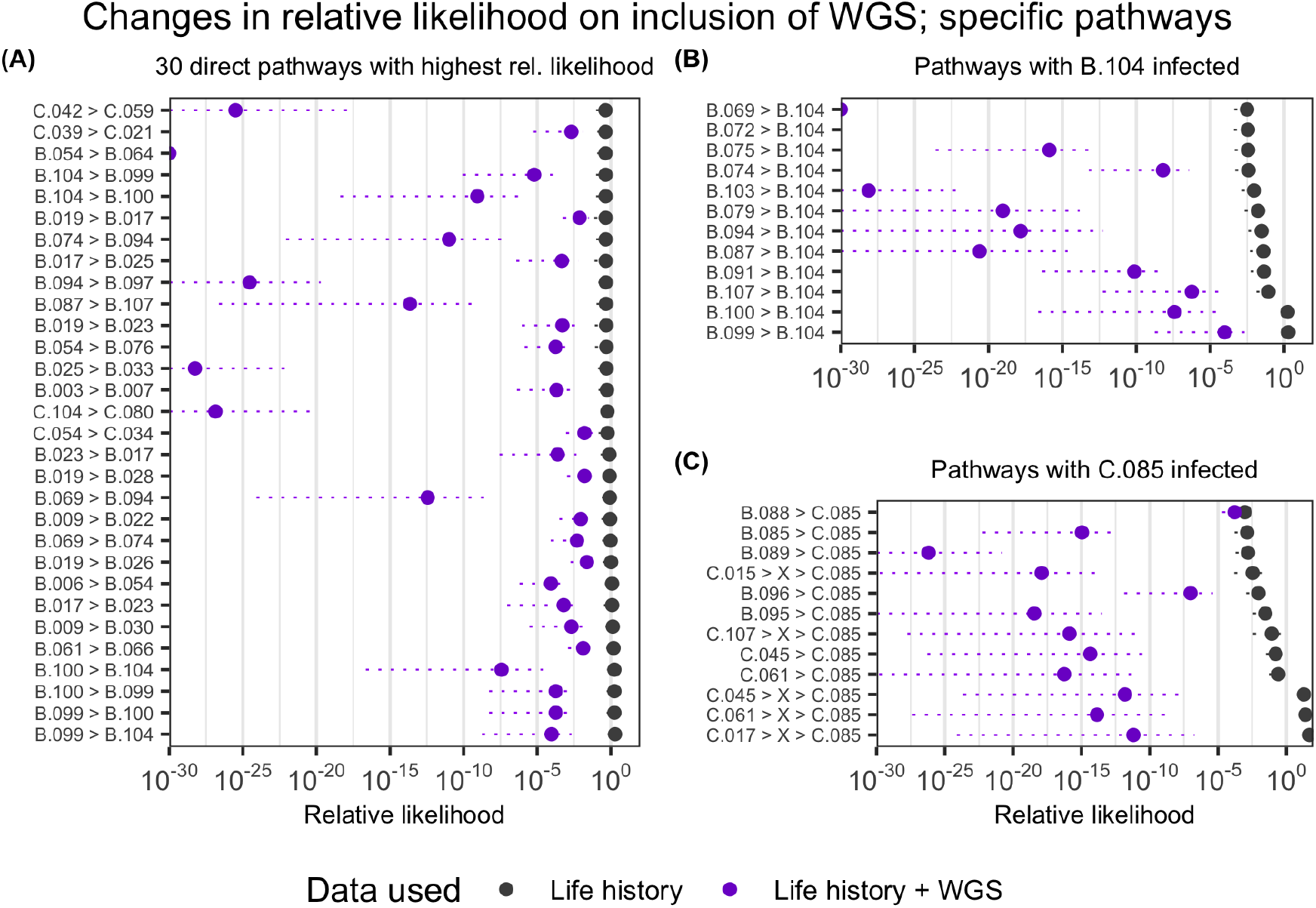
(A): Differences in relative likelihoods for the 30 pathways estimated as having the highest relative likelihood based on life history data alone, on the inclusion of WGS data. The drop in relative likelihood differs by several orders of magnitude across different pathways. The dotted lines indicate the range of the [0.05, 0.95] confidence interval of the relative likelihood evaluated over different parameter samples. (B), (C): the 12 highest relative likelihood transmission pathways where hosts B.104 and C.085 are the infectees, respectively.

## F Sensitivity of relative likelihood to parameter changes

**Figure S9:**
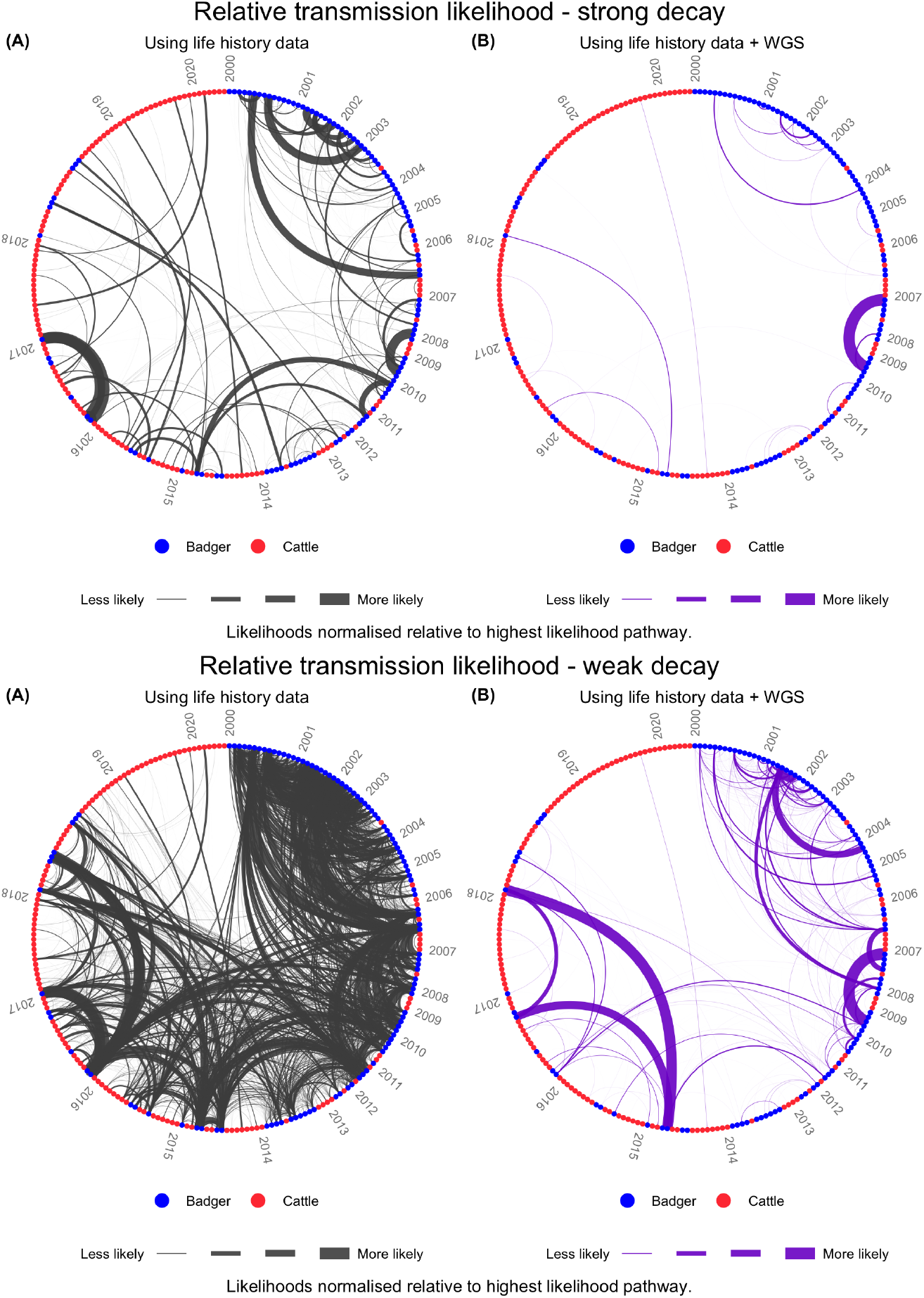
Relative transmission likelihoods over all direct pathways as per Figure 1, but with the exponential rate of decay in the infection rate *α* being 1=3 times the value used in the main analysis (top), and 3 times its value (bottom).

**Figure S10:**
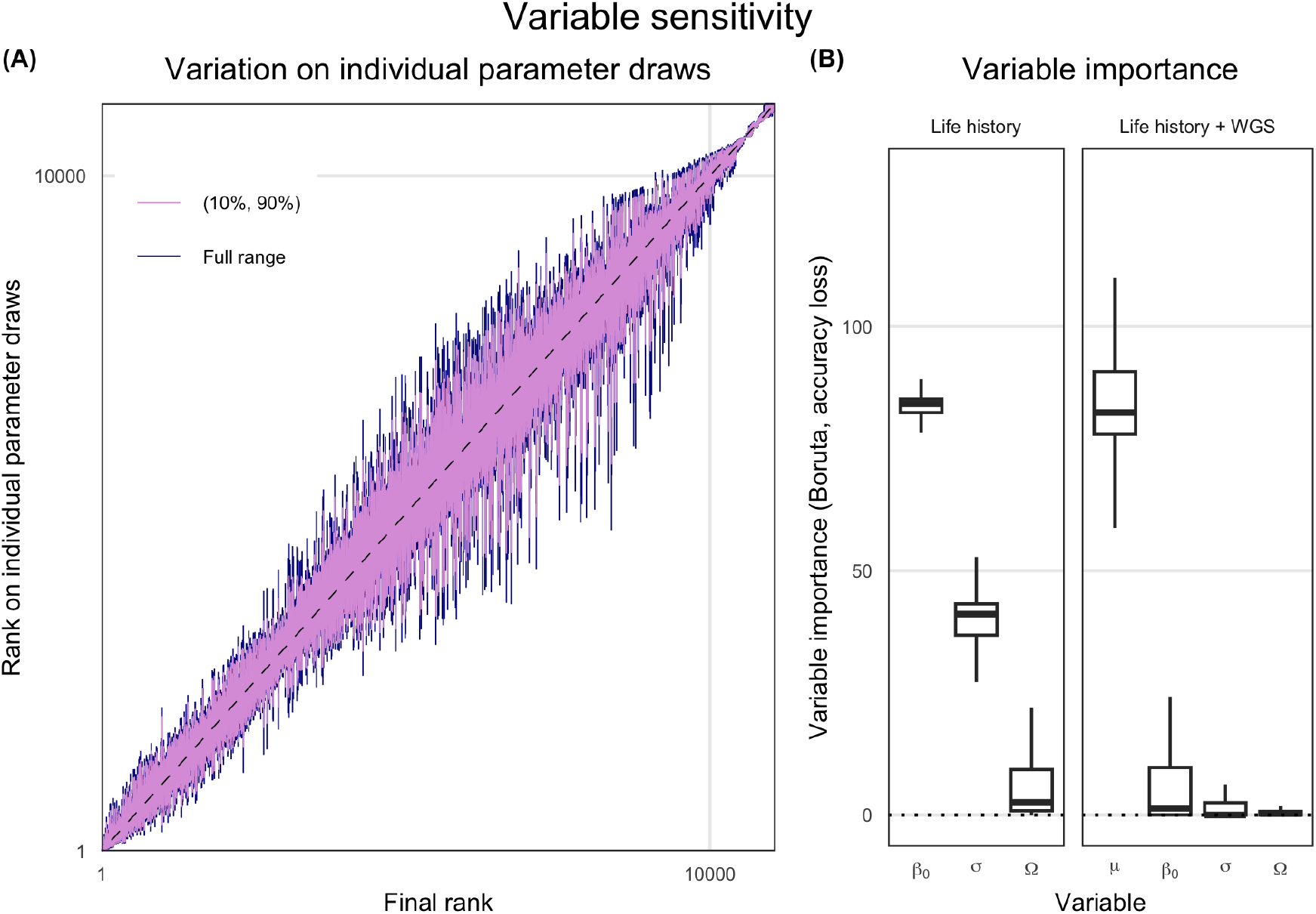
Sensitivity analyses. (**A**) the variability of each transmission pathway’s rank over di_erent parameter draws. Pathways are ordered on the *x*-axis in order of final rank (1 = highest relative likelihood), and the range of ranks over different parameter draws are shown on the *y*-axis. Pathways with median likelihood of zero are excluded. (**B**) Results of applying a Boruta algorithm to the relative likelihoods, assessing the relative importance of different parameters in influencing the value of the relative likelihood. The box plot indicates the distribution of importances over different pathways.

## References

[1] Pybus OG, Rambaut A. Evolutionary analysis of the dynamics of viral infectious disease. Nature Reviews Genetics. 2009;10(8):540–550.

[2] Kao RR, Haydon DT, Lycett SJ, Murcia PR. Supersize me: how whole-genome sequencing and big data are transforming epidemiology. Trends in microbiology. 2014;22(5):282–291.

[3] Ladner JT, Grubaugh ND, Pybus OG, Andersen KG. Precision epidemiology for infectious disease control. Nature medicine. 2019;25(2):206–211.

[4] Guinat C, Vergne T, Kocher A, Chakraborty D, Paul MC, Ducatez M, et al. What can phylodynamics bring to animal health research? Trends in Ecology & Evolution. 2021;36(9):837– 847.

[5] Bromham L, Penny D. The modern molecular clock. Nature Reviews Genetics. 2003;4(3):216–224.

[6] Menardo F, Duchene S, Brites D, Gagneux S. The molecular clock of Mycobacterium tuberculosis. PLoS pathogens. 2019;15(9):e1008067.

[7] Sanjuan R, Nebot MR, Chirico N, Mansky LM, Belshaw R. Viral mutation rates. Journal of virology. 2010;84(19):9733–9748.

[8] Biek R, Pybus OG, Lloyd-Smith JO, Didelot X. Measurably evolving pathogens in the genomic era. Trends in ecology & evolution. 2015;30(6):306–313.

[9] O’Reilly LM, Daborn C. The epidemiology of Mycobacterium bovis infections in animals and man: a review. Tubercle and Lung disease. 1995;76:1–46.

[10] Garnier T, Eiglmeier K, Camus JC, Medina N, Mansoor H, Pryor M, et al. The complete genome sequence of Mycobacterium bovis. Proceedings of the National Academy of Sciences. 2003;100(13):7877–7882.

[11] Crispell J, Zadoks RN, Harris SR, Paterson B, Collins DM, de Lisle GW, et al. Using whole genome sequencing to investigate transmission in a multi-host system: bovine tuberculosis in New Zealand. BMC genomics. 2017;18:1–12.

[12] Crispell J, Benton CH, Balaz D, De Maio N, Ahkmetova A, Allen A, et al. Combining genomics and epidemiology to analyse bi-directional transmission of Mycobacterium bovis in a multi-host system. Elife. 2019;8:e45833.

[13] Trewby H, Wright D, Breadon EL, Lycett SJ, Mallon TR, McCormick C, et al. Use of bacterial whole-genome sequencing to investigate local persistence and spread in bovine tuberculosis. Epidemics. 2016;14:26–35.

[14] Akhmetova A, Guerrero J, McAdam P, Salvador LC, Crispell J, Lavery J, et al. Genomic epidemiology of Mycobacterium bovis infection in sympatric badger and cattle populations in Northern Ireland. Microbial Genomics. 2023;9(5).

[15] Canini L, Modenesi G, Courcoul A, Boschiroli ML, Durand B, Michelet L. Deciphering the role of host species for two Mycobacterium bovis genotypes from the European 3 clonal complex circulation within a cattle-badger-wild boar multihost system. MicrobiologyOpen. 2023;12(1):e1331.

[16] O’Brien DJ, Kao RR, Little RA, Enticott G, Riley SJ. The Road Not Traveled: Bovine Tuberculosis in England, Wales, and Michigan, USA. One Health Cases. 2023;.

[17] Butler AJ, Lobley M, Winter M. Economic impact assessment of bovine tuberculosis in the south west of England; 2010.

[18] Barnes AP, Moxey A, Brocklehurst S, Barratt A, McKendrick IJ, Innocent G, et al. The consequential costs of bovine tuberculosis (bTB) breakdowns in England and Wales. Preventive Veterinary Medicine. 2023;211:105808.

[19] De la Rua-Domenech R, Goodchild A, Vordermeier H, Hewinson R, Christiansen K, Clifton-Hadley R. Ante mortem diagnosis of tuberculosis in cattle: a review of the tuberculin tests, γ-interferon assay and other ancillary diagnostic techniques. Research in veterinary science. 2006;81(2):190–210.

[20] van Tonder AJ, Thornton MJ, Conlan AJ, Jolley KA, Goolding L, Mitchell AP, et al. Inferring Mycobacterium bovis transmission between cattle and badgers using isolates from the Randomised Badger Culling Trial. PLoS Pathogens. 2021;17(11):e1010075.

[21] Reis AC, Salvador LC, Robbe-Austerman S, Tenreiro R, Botelho A, Albuquerque T, et al. Whole genome sequencing refines knowledge on the population structure of Mycobacterium bovis from a multi-host tuberculosis system. Microorganisms. 2021;9(8):1585.

[22] Perea C, Ciaravino G, Stuber T, Thacker TC, Robbe-Austerman S, Allepuz A, et al. Wholegenome SNP analysis identifies putative Mycobacterium bovis transmission clusters in live-stock and wildlife in Catalonia, Spain. Microorganisms. 2021;9(8):1629.

[23] Rodrigues RdA, Ribeiro Araujo F, Rivera Davila AM, Etges RN, Parkhill J, van Tonder AJ. Genomic and temporal analyses of Mycobacterium bovis in southern Brazil. Microbial Genomics. 2021;7(5):000569.

[24] Rossi G, Shih BBJ, Egbe NF, Motta P, Duchatel F, Kelly RF, et al. Unraveling the epidemiology of Mycobacterium bovis using whole-genome sequencing combined with environmental and demographic data. Frontiers in Veterinary Science. 2023;10:1086001.

[25] Biek R, O’Hare A, Wright D, Mallon T, McCormick C, Orton RJ, et al. Whole genome sequencing reveals local transmission patterns of Mycobacterium bovis in sympatric cattle and badger populations. PLoS pathogens. 2012;8(11):e1003008.

[26] Santos N, Colino EF, Arnal MC, de Luco DF, Sevilla I, Garrido JM, et al. Complementary roles of wild boar and red deer to animal tuberculosis maintenance in multi-host communities. Epidemics. 2022;41:100633.

[27] Price-Carter M, Brauning R, De Lisle GW, Livingstone P, Neill M, Sinclair J, et al. Whole genome sequencing for determining the source of Mycobacterium bovis infections in livestock herds and wildlife in New Zealand. Frontiers in Veterinary Science. 2018;5:272.

[28] Rossi G, Crispell J, Balaz D, Lycett SJ, Benton CH, Delahay RJ, et al. Identifying likely transmissions in Mycobacterium bovis infected populations of cattle and badgers using the Kolmogorov Forward Equations. Scientific Reports. 2020;10(1):21980.

[29] Rossi G, Crispell J, Brough T, Lycett SJ, White PC, Allen A, et al. Phylodynamic analysis of an emergent Mycobacterium bovis outbreak in an area with no previously known wildlife infections. Journal of Applied Ecology. 2022;59(1):210–222.

[30] Pereira AC, Reis AC, Cunha MV. Genomic epidemiology sheds light on the emergence and spread of Mycobacterium bovis Eu2 Clonal Complex in Portugal. Emerging Microbes & Infections. 2023;12(2):2253340.

[31] Campbell F, Strang C, Ferguson N, Cori A, Jombart T. When are pathogen genome sequences informative of transmission events? PLoS pathogens. 2018;14(2):e1006885.

[32] Duault H, Durand B, Canini L. Methods combining genomic and epidemiological data in the reconstruction of transmission trees: a systematic review. Pathogens. 2022;11(2):252.

[33] Didelot X, Fraser C, Gardy J, Colijn C. Genomic infectious disease epidemiology in partially sampled and ongoing outbreaks. Molecular biology and evolution. 2017;34(4):997–1007.

[34] Xu Y, Cancino-Munoz I, Torres-Puente M, Villamayor LM, Borras R, Borras-Manez M, et al. High-resolution mapping of tuberculosis transmission: Whole genome sequencing and phylogenetic modelling of a cohort from Valencia Region, Spain. PLoS medicine. 2019;16(10):e1002961.

[35] Sobkowiak B, Banda L, Mzembe T, Crampin AC, Glynn JR, Clark TG. Bayesian reconstruction of Mycobacterium tuberculosis transmission networks in a high incidence area over two decades in Malawi reveals associated risk factors and genomic variants. Microbial genomics. 2020;6(4):e000361.

[36] Xu Y, Stockdale JE, Naidu V, Hatherell H, Stimson J, Stagg HR, et al. Transmission analysis of a large tuberculosis outbreak in London: a mathematical modelling study using genomic data. Microbial genomics. 2020;6(11):e000450.

[37] Sobkowiak B, Romanowski K, Sekirov I, Gardy JL, Johnston JC. Comparing Mycobacterium tuberculosis transmission reconstruction models from whole genome sequence data. Epidemiology & Infection. 2023;151:e105.

[38] Duault H, Durand B, Canini L. Outbreak reconstruction with a slowly evolving multi-host pathogen: a comparative study of three existing methods on Mycobacterium bovis outbreaks. bioRxiv. 2023; p. 2023–07.

[39] Godfray HCJ, Donnelly CA, Kao RR, Macdonald DW, McDonald RA, Petrokofsky G, et al. A restatement of the natural science evidence base relevant to the control of bovine tuberculosis in Great Britain. Proceedings of the Royal Society B: Biological Sciences. 2013;280(1768):20131634.

[40] Delahay RJ, Walker N, Smith G, Wilkinson D, Clifton-Hadley R, Cheeseman C, et al. Longterm temporal trends and estimated transmission rates for Mycobacterium bovis infection in an undisturbed high-density badger (Meles meles) population. Epidemiology & Infection. 2013;141(7):1445–1456.

[41] Gaughran A, Kelly DJ, MacWhite T, Mullen E, Maher P, Good M, et al. Super-ranging. A new ranging strategy in European badgers. PLoS One. 2018;13(2):e0191818.

[42] Nunez-Garcia J, Downs SH, Parry JE, Abernethy DA, Broughan JM, Cameron AR, et al. Meta-analyses of the sensitivity and specificity of ante-mortem and post-mortem diagnostic tests for bovine tuberculosis in the UK and Ireland. Preventive Veterinary Medicine. 2018;153:94–107.

[43] Animal and Plant Health Agency. Guidance — Bovine TB testing intervals; 2020. [Online; accessed 17 November 2023]. https://www.gov.uk/guidance/bovine-tb-testing-intervals.

[44] Animal and Plant Health Agency. APHA Briefing Note 13/21; Introduction of the Operational Use of Whole Genome Sequencing (WGS) for bTB Purposes; 2021. [Online; accessed 9 November 2023]. http://apha.defra.gov.uk/documents/ov/Briefing-Note-1321.pdf.

[45] Animal and Plant Health Agency. Bovine tuberculosis in Great Britain in 2022; Explanatory Supplement to the annual reports; 2022. [Online; accessed 9 November 2023]. BovinetuberculosisinGreatBritainin2022ExplanatorySupplementtotheannualreports.

[46] Malone KM, Farrell D, Stuber TP, Schubert OT, Aebersold R, Robbe-Austerman S, et al. Updated reference genome sequence and annotation of Mycobacterium bovis AF2122/97. Genome announcements. 2017;5(14):10–1128.

[47] Delahay R, Smith G, Ward A, Cl C. Options for the management of bovine tuberculosis transmission from badger (Meles meles) to cattle: evidence from a long-term study. Mammal Study. 2005 12;30:S73–S81.

[48] McDonald JL, Robertson A, Silk MJ. Wildlife disease ecology from the individual to the population: Insights from a long-term study of a naturally infected European badger population. Journal of Animal Ecology. 2018;87(1):101–112.

[49] Department for Environment, Food & Rural Affairs. CTS Online;. [Online; accessed 10 November 2023]. https://secure.services.defra.gov.uk/wps/portal/ctso.

[50] Animal and Plant Health Agency. SAM;. [Online; accessed 10 November 2023]. https://www.data.gov.uk/dataset/a553b50e-60ab-45f7-9b6c-90d0c21ada80/sam.

[51] Cheeseman C. Badgers. T & AD Poyser Natural History; 1996.

[52] Conlan AJ, McKinley TJ, Karolemeas K, Pollock EB, Goodchild AV, Mitchell AP, et al. Estimating the hidden burden of bovine tuberculosis in Great Britain. 2012;.

[53] Dean GS, Rhodes SG, Coad M, Whelan AO, Cockle PJ, Clifford DJ, et al. Minimum infective dose of Mycobacterium bovis in cattle. Infection and immunity. 2005;73(10):6467–6471.

[54] Brooks-Pollock E, Roberts GO, Keeling MJ. A dynamic model of bovine tuberculosis spread and control in Great Britain. Nature. 2014;511(7508):228–231.

[55] Rogers L, Cheeseman C, Mallinson P, Clifton-Hadley R. The demography of a high-density badger (Meles meles) population in the west of England. Journal of Zoology. 1997;242(4):705– 728.

[56] Vicente J, Delahay R, Walker N, Cheeseman C. Social organization and movement influence the incidence of bovine tuberculosis in an undisturbed high-density badger Meles meles population. Journal of Animal Ecology. 2007;76(2):348–360.

[57] Rogers L, Delahay R, Cheeseman C, Langton S, Smith G, Clifton-Hadley R. Movement of badgers (Meles meles) in a high–density population: individual, population and dis-ease effects. Proceedings of the Royal Society of London Series B: Biological Sciences. 1998;265(1403):1269–1276.

[58] Dugdale HL, Macdonald DW, Pope LC, Johnson PJ, Burke T. Reproductive skew and relatedness in social groups of European badgers, Meles meles. Molecular Ecology. 2008;17(7):1815– 1827.

[59] Benton CH, Delahay RJ, Smith FA, Robertson A, McDonald RA, Young AJ, et al. Inbreeding intensifies sex-and age-dependent disease in a wild mammal. Journal of Animal Ecology. 2018;87(6):1500–1511.

[60] Bohm M, Palphramand KL, Newton-Cross G, Hutchings MR, White PC. Dynamic interactions among badgers: implications for sociality and disease transmission. Journal of Animal Ecology. 2008;77(4):735–745.

[61] Macdonald DW, Newman C, Buesching CD, Johnson PJ. Male-biased movement in a high-density population of the Eurasian badger (Meles meles). Journal of Mammalogy. 2008;89(5):1077–1086.

[62] Smith G, Cheeseman C, Wilkinson D, Clifton-Hadley R. A model of bovine tuberculosis in the badger Meles meles: the inclusion of cattle and the use of a live test. Journal of Applied Ecology. 2001; p. 520–535.

[63] R Core Team. R: A Language and Environment for Statistical Computing. Vienna, Austria; 2022. Available from: https://www.R-project.org/.

[64] Wolfram Research, Inc. Mathematica, Version 12.3.1.0;. Champaign, IL, 2023. Available from: https://www.wolfram.com/mathematica.

